# The Importance of Nonsense Errors: Estimating the Rate and Implications of Drop-Off Errors during Protein Synthesis

**DOI:** 10.1101/2024.09.05.611510

**Authors:** Alexander L. Cope, Denizhan Pak, Michael A. Gilchrist

## Abstract

The process of mRNA translation is both energetically costly and relatively error-prone compared to transcription and replication. Nonsense errors during mRNA translation occur when a ribosome drops off a transcript before reaching a stop codon, resulting in energetic investment in an incomplete and likely non-functional protein. Nonsense errors impose a potentially significant energy burden on the cell, making it critical to quantify their frequency and energetic cost. Here, we present a model of ribosome movement for estimating protein production, elongation, and nonsense error rates from high-throughput ribosome profiling data. Applying this model to an exemplary ribosome profiling dataset in *S. cerevisiae*, we find that nonsense error rates vary between codons, in conflict with the general assumption of uniform rates across sense codons. Using our parameter estimates, we find multiple lines of evidence that selection against nonsense errors is a prominent force shaping coding-sequence evolution, including that nonsense errors place an energetic burden on cells comparable to ribosome pausing. Our results indicate greater consideration should be given to the impact of nonsense errors in shaping coding-sequence evolution.

**Author Summary:** The process of mRNA translation is both energetically expensive and relatively error-prone. As such, natural selection is thought to shape the evolution of CDSs to reduce the cost of these errors when they occur. Nonsense errors (NSEs) occur when a ribosome stops translation prior to completing a functional protein, resulting in wasted energy on non-functional product. Despite their functional consequences, NSEs and their effects on sequence evolution are generally understudied compared to other types of translation errors. This is in part due to the challenge of quantifying these errors from omics-scale data. We present a model for quantifying codon-specific estimates of elongation and NSE rates from ribosome profiling data, which gives a snapshot of the actively translating ribosomes in a cell. Although it is well-established that sense codons vary in their elongation rates, we find evidence that codons also vary in their NSE rates. Using our parameter estimates, we find multiple lines of evidence for selection against NSEs shaping patterns of codon usage bias. Our results suggest the cost of NSEs are comparable to the cost of ribosome pausing, and thus may play a greater role in coding-sequence evolution than previously appreciated.

## Introduction

Protein synthesis, i.e., the transcription and translation of mRNAs, is one of the most energetically costly cellular processes [1]. For example, 66% of ATP is dedicated to protein synthesis in *E. coli* [2]. While both transcription and translation are critical to protein synthesis, the cost of translation is estimated to be 125 times the cost of transcription [3]. Translation involves both direct (GTPs used to charge tRNAs and drive each elongation cycle) and indirect costs (overhead cost of the translation infrastructure such as the ribosome and tRNAs) [4–6]. Furthermore, mRNAs generally undergo multiple rounds of translation. Given the energetic cost of mRNA translation and the finite energy budget of a cell, genotypes reducing the cost of protein production are expected to have a selective advantage that scales with a gene’s average protein production rate. A classic example of this is the bias of highly expressed genes toward synonymous codons recognized by more abundant tRNA, which is hypothesized to be an adaptation due to natural selection to reduce the indirect cost of ribosome pausing [5–8].

In addition to being energetically costly, mRNA translation is relatively error-prone compared to DNA replication and transcription [9]. Translation errors potentially result in non-functional proteins. Much of the literature has focused on the impact of codons being misread by the ribosome, resulting in the insertion of the incorrect amino acid into the growing peptide chain (i.e., a missense error) [10–13]. The incorrect amino acid sequence may result in a misfolded protein that is potentially corrected via interactions with chaperone proteins [14, 15]. In contrast, a nonsense error (a.k.a. premature termination errors or processivity errors) results in an incomplete protein [16, 17]. As an incomplete protein is expected to have little if any functional capacity and is unlikely to be rescued via chaperones, nonsense errors may contribute substantially to the overall cost of mRNA translation [18]. Based on work in *E. coli* and *S. cerevisiae*, nonsense errors are estimated to occur at a frequency of approximately 1 every 10,000 codons [19–23].

Each ribosome elongation step can be thought of as having two possible outcomes: a nonsense error or successful elongation. Assuming the amount of time a ribosome spends waiting before elongation or an error occurs is exponentially distributed, the probability of a nonsense error at a given codon depends on both its nonsense error rate (i.e., the background rate at which a nonsense error event is triggered) and its elongation rate. It is well-established that ribosome elongation rates generally vary across codons, often due to differences in the tRNA availability and codon-anticodon (e.g., wobble) base-pairing effects [24–29]. As a result, slower codons are expected to be more prone to nonsense errors even if nonsense error rates are uniform across codons [18, 30–32].

Many theoretical studies of mRNA translation dynamics assume the nonsense error rate is uniform across codons. Theoretical studies making this assumption, while still allowing variable ribosome elongation rates across codons, predict the probability of a ribosome completing translation varies substantially across genes as a function of amino acid sequence, protein length, and codon usage bias [30, 33, 34]. Conflicting with the assumption of uniformity, studies in both *E. coli* and *S. cerevisiae* indicate sense codons differ in their capacity to pair with release factors [35, 36], often by mimicking the structural mechanism of stop codon recognition upon release factor binding [37]. Additionally, other mechanisms that lead to nonsense errors, such as peptidyl-tRNA drop-off [38–40] and ribosomal frameshifting [41, 42], are expected to vary across codons. This limited evidence suggests nonsense error rates differ across codons, which could amplify or dampen the differences in codon-specific elongation probabilities. If codons vary in their nonsense error probabilities, then it follows that protein-coding DNA sequences (from here on out, CDS) can evolve to reduce the cost of nonsense errors via codon usage. As before, the selective advantage for synonymous genotypes reducing the cost of nonsense errors should increase with a gene’s average protein production rate. Thus, we expect to see increasing evidence of adaptation in codon usage bias to reduce the cost of protein production with a gene’s average protein production rate.

Ribosome profiling is a powerful technique for quantifying steady-state translation across the transcriptome [43]. Compared to mass spectrometry-based approaches (including those specifically intended to quantify translation), ribosome profiling provides better sequence coverage, a broader dynamic range, and reveals actively translating ribosomes at codon-level resolution [44]. Due to these advantages and the ever decreasing monetary cost of next-generation sequencing technologies, ribosome profiling is a powerful approach for studying translation on an omics-scale [45]. However, ribosome profiling does not directly track ribosome movement along a transcript, requiring mathematical models to extract biological information from these empirical measurements. Previous attempts to quantify nonsense errors from ribosome profiling data have typically averaged over codons and genes, ignoring variation in elongation, nonsense, and protein production rates [21, 23]. To explicitly account for this variation, we extend our previous model of nonsense errors and ribosome movement along an mRNA transcript [18, 30] to estimate codon-specific elongation and nonsense error rates using a high-quality ribosome profiling dataset in *S. cerevisiae* [27].

Using our estimates, we evaluate and contextualize the estimated distribution and costs of nonsense errors across the *S. cerevisiae* transcriptome. We find compelling evidence that nonsense error rates and probabilities differ across codons and amino acids. As genes differ in amino acid usage, protein length, and codon usage, our results indicate the probability a ribosome completes translation varies across genes (interquartile range 0.87 – 0.95). We identify multiple lines of evidence that the yeast genome is extensively adapted to reduce the cost of nonsense errors. Our evidence includes a bias towards nonsense error-prone codons near the 5’-ends of transcripts and an avoidance of nonsense error-prone codons in highly expressed genes. Consistent with previous theoretical studies relying upon tRNA-based proxies of elongation rates [30, 33], we find approximately 60% of genes in *S. cerevisiae* exhibit signals of adaptation to reduce the cost of nonsense errors. Despite these adaptions to reduce the cost of nonsense errors, we estimate nonsense errors impose an energetic burden comparable to, if not greater than, ribosome pausing. Although nonsense errors are believed to have a lower probability than missense errors, because a preponderance of nonsense errors (but only a fraction of missense errors) are likely to disrupt a protein’s functionality, our work suggests natural selection against nonsense errors plays a substantial, important, and underappreciated role in coding sequence evolution.

## Materials and Methods

We are interested in calculating the probability of observing a ribosome footprint (RFP i.e., a mapped sequencing read) at a codon within a transcript as measured via ribosome profiling. We assume there is a pool of RFP generated from the transcriptome and that the mRNAs in this pool are close to steady-state in terms of ribosome initiation and completion of translating a transcript. Below we give an overview of our model formulation and usage, with additional details found in S1 Text. Definitions of all model parameters can be found in Table 1.

**Table 1:** Table of model parameters.

| Parameter | Definition | Units |
| --- | --- | --- |
| $c_{g,i}$ | Ribosome elongation rate at codon position $i$ in gene $g$ | 1/t |
| $b$ | Ribosome nonsense error rate for codon $c$ | 1/t |
| $\alpha_c, \lambda_c$ | Shape and scale parameters for Inverse Gamma distribution of codon $c$ elongation rates. | |
| $m_g$ | Density of mRNA transcripts for gene $g$ in cytosol. | 1/Vol |
| $M_g$ | Observed mRNA counts from RNA-Seq data. | 1/Vol |
| $\kappa_g$ | Rate constant determining ribosome initiation rate per mRNA. Function of diffusion of ribosomes, mRNA, and other factors. | 1/t |
| $\kappa'_g$ | Rate of initiation of protein synthesis under a given set of conditions. Equal to $m_g \kappa_g$. | 1/t |
| $\sigma_g(i-1)$ | Probability ribosome reaches codon $i$ by successfully translating the previous $i-1$ codons of codon $g$. | |
| $\mathbb{E}[\sigma_g(i-1)]$ | Expectation of $\sigma_g(i-1)$ used when fitting model to data. | |
| $p_{g,i}$ | Probability a ribosome is found at position $i$ on an mRNA transcript from gene $g$ when translation initiation and completion of mRNA is at steady state. | |
| $P_{g,i}$ | Probability of observing a footprint for position $i$ of mRNA from gene $g$. | |
| $P_g^c$ | Probability of observing a footprint for codon $c$ of mRNA from gene $g$. | |
| $Z$ | Partition function which scales the codon footprint sampling probabilities $P_{g,i}$ summed across $i$ and $g$ equals 1. | |
| $n_g$ | Number of codons in mRNA of gene $g$. | |
| $Y_{g,i}$ | Number of RFP observed for position $i$ in gene $g$ | |
| $Y$ | Total number of RFP in dataset. | |
| $\lambda'_c$ | Composite parameter equal to $\lambda_c Z / Y$ | |
| $\pi_j$ | Prior probability for parameter $j$. | |

### Ribosome Pausing and Nonsense Error Model

Below we outline the assumptions behind our ribosome PAusing and NonSense Error (PANSE) model and the formal likelihood function they lead to.

### PANSE Model Assumptions

For a given gene *g, m*_*g*_ represents its equilibrium mRNA transcript concentration within a cell and *κ*_*g*_ represents the average ribosome initiation rate for each of its mRNAs. Gene *g* consists of *n*_*g*_ codons and the elongation and nonsense error rates at codon position *i* are *c*_*g,i*_ and *b*_*g,i*_, respectively. We assume the elongation rate *c*_*g,i*_ varies between sites such that *c*_*g,i*_ ∼ InverseGamma(*α*_*i*_, *λ*_*i*_) where the shape parameter *α*_*i*_ and scale parameter *λ*_*i*_ can vary between the 61 sense codons. The across-site heterogeneity of a given sense codon’s elongation rates reflects a variety of factors that alter ribosome elongation [22, 46]. Although the nonsense error rate *b*_*g,i*_ likely varies with codon and position, for simplicity we focus solely on the variation between codons and, thus, treat the codon-specific nonsense error rate *b* as a constant, having the same value across all codons of type *c*. Based on these assumptions, it is possible to model the distribution of footprints as coming from a negative binomial distribution, i.e. *Y*_*g,i*_ ∼ NB(*k, p*) with 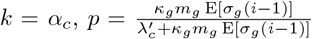, where 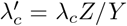. The composite parameter 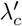 is a rescaling of the codons specific rate parameter *λ*_*c*_ by the ratio of the partition function for the RFP distribution *Z* the footprints are sampled from to the total number of observed footprints in a dataset *Y* = ∑_*g,i*_ *Y*_*g,i*_. Thus, Z/*Y* is a measure of the sampling efficiency of the experiment.

Given *Y*_*g,i*_ represents the observed number of ribosome footprints (RFP) at codon *i* of gene *g*, it follows that the likelihood of observing *Y*_*g,i*_ is,

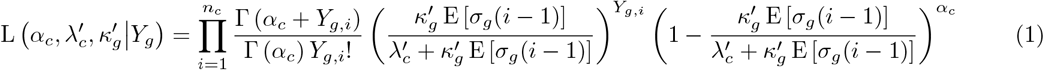

where 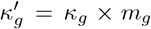 is a rescaling of the mRNA_*g*_ specific ribosome initiation rate *κ*_*g*_ by the density of mRNA_*g*_ transcripts within the cell, *m*_*g*_. Assuming independence between elongation steps, it follows that the probability of a ribosome successfully elongating at codons 1 through *i* − 1 is,

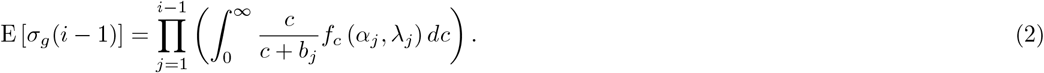

where *f*_*c*_(*α*_*i*_, *λ*_*i*_) represents the PDF of the InverseGamma distribution for the ribosome elongation rate for codon *i*.

Because the nonsense error rate *b* between sense codons is likely to be several orders of magnitude less than a given *c*_*g,i*_ and we will be working with our likelihood function on the log scale, we can greatly speed up our evaluation of the log of Eqn. (1) by approximating log (E [*σ*_*g*_(*i*)]) using a 2nd order Taylor Series approximation around a mean nonsense error rate for all codons 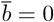. Doing so gives,

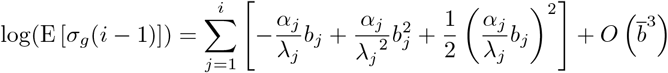

### Analysis of ribosome profiling data

Raw ribosome profiling for *S. cerevisiae* reads were downloaded from the Sequence Read Archive (SRA Run Accession: SRR1049521) [27]. These data were generated using a flash-freeze protocol to halt ribosome elongation, which avoids many of the technical artifacts caused by the use of cycloheximide [27, 47]. The raw reads were processed using the riboviz2 [48]. Aligned reads of length 28-30 were extracted and assigned to codons in the A-site assuming a 15 nucleotide offset relative to the 5’-end of the read [43] (S1 Fig). We note that using different A-site offsets based on the riboWaltz R package had little impact on parameter estimation for the Weinberg et al. data (S2 Fig).

A flat file was created that includes the number of ribosome footprints (i.e., counts) at each codon within a gene. This file was used as the primary input into our implementation of PANSE. We employed numerous filtering strategies to remove genes that may violate the assumptions of the model (see Supplementary Material: Filtering of processed ribosome profiling data and S3 Fig). This ultimately led to a final dataset with 3,112 genes, approximately 50% of the CDSs in *S. cerevisiae*.

### Fitting the Pausing and Nonsense Error Model

The PANSE model was fit via a Markov chain Monte Carlo (MCMC) algorithm to the codon-level ribosome footprint counts across the approximately 3,000 genes included in the final data set. The MCMC was run for 50,0000 iterations, keeping every 5th iteration to reduce autocorrelation. Convergence was assessed by comparing the results of two separate MCMCs that were started at random points in parameter space. Posterior means and 95% highest density intervals (HDIs) were calculated for each parameter of PANSE based on the MCMC traces. Consistent with our previous work [7], gene-specific initiation rates 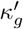 are assumed to follow a lognormal distribution with a mean initiation rate of 1. This is accomplished by fixing the mean of the lognormal distribution to be 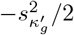, where 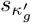 is the standard deviation of the lognormal distribution. The shape and scale parameters, *α* and *λ*, for each codon specific elongation rate, *c*, and the codon specific nonsense error rates *b*, were assumed to have a flat prior with ranges (0, 100), (0, 100) and (1e-100, 1e-1) on the natural scale. We note these are particularly broad distributions, but we wanted to ensure that adequate parameter space could be explored. We fit PANSE (1) assuming no NSEs occur, (2) assuming uniform NSE rates across codons and (3) allowing NSE rates to vary across codons. To determine if there is support for variation in NSE rates across codons, these three model fits were compared using the Deviance Information Criterion (DIC) [49].

The ribosome profiling data we used were prepared with a flash-freeze protocol to halt elongation, but there is still an observable increase in ribosome density in the first 200 codons of the transcript. Although the increased density was less so than observed in ribosome profiling measurements made with cycloheximide, it remains unclear if this increased density reflects true biology or a technical artifact [27]. As it is plausible that ribosome counts for the first 200 codons are impacted by unknown technical biases, we fit PANSE masking the ribosome counts for this region in the likelihood calculation for each gene. However, we did include these codons in our calculation of the expected probability of elongation up to codon *i E*[*σ*_*g*_(*i*)]. We compared parameter estimates of PANSE to (1) independent empirical data such as mRNA abundances and tRNA-based proxies for elongation rates [27], and (2) parameter estimates from the Ribosomal Overhead Cost version of the Stochastic Evolutionary Model of Protein Production Rates (ROC-SEMPPR), which estimates protein production rates *ϕ* and codon-specific selection coefficients Δ*η* from codon usage bias patterns [7]. See previous work regarding parameter estimation with ROC-SEMPPR for *S. cerevisiae* [7, 50].

### Analyzing variation in nonsense error rates *b* across codons

To better understand the variation in nonsense error (NSE) rates across codons, we performed a linear regression of the posterior mean NSE rates *b* against numerous properties of the codon and amino acid. This includes the identity of the nucleotide at each of the three codon positions (A, G, C, or T, with A as the reference class), the number of stop codons 1 nucleotide mutation away from the codon (0, 1, or 2), and the physicochemical properties of the resulting amino acid (charged, hydrophobic, neutral, other) on each codon. We weighted each codon in the regression by the standard deviations estimated from the MCMC traces.

### Quantifying variation in translation completion probabilities *σ*_*g*_(*n*_*g*_) across genes

We used our estimates of elongation and NSE rates, *c* and *b*, for each of the 61 sense codons to calculate the translation completion probability *σ*_*g*_(*n*_*g*_). This allowed us to apply this formula to all genes in the *S. cerevisiae* genome, regardless of whether or not it was included in the model fit. We assessed how variation in translation completion probabilities *σ*_*g*_(*n*_*g*_) varied across genes as a function of length and gene expression measured as mRNA abundances (in units of RPKM) taken from previous work [27]. Additionally, we compared how well our inferences of translation completion probabilities *σ*_*g*_(*n*_*g*_) made directly from empirical ribosome profiling data compared to theoretical estimates based on simulations from a Totally Asymmetric Exclusion Process (TASEP) model of translation [34].

### Quantifying elongation probability variation within and across genes

As before, *c*_*i*_ and *b*_*i*_ represent the expected elongation and nonsense error rates the codon of at position *i*, respectively. Given that there are only two possible outcomes in our model (elongation or a nonsense error), it follows that the probability a ribosome elongates the growing peptide chain at codon *i* is,

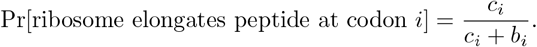

The probability a ribosome reaches codon *i* by successfully elongating at the previous *i* − 1 codons is,

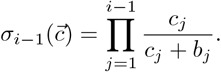

To evaluate how elongation probabilities vary with codon position *i*, we calculated the average probability a ribosome elongates at a given position as the geometric mean of the elongation probability across genes. We then regressed the log(Position *i*) against the average elongation probability to quantify how the elongation probability changes as a function of position. To test if the observed slope was greater than expected under the null model of no selection against nonsense errors, we generated 1000 different permuted sets of genes for each of 3 different possible nulls: (1) the synonymous codons of an amino acid were permuted within a gene (randomized by amino acid), (2) amino acids and codons were permuted within a gene (randomized by CDS), and (3) codons were permuted across genes. The slope estimated from the real set of genes was then compared to each of the 3 null distributions, with the *p*-value= (*k* + 1)*/*(1000 + 1), where *k* is the number of occurrences in which a slope from a permuted set of genes was greater than the slope from the real set of genes. A similar analysis was performed on the real data after binning genes based on mRNA abundances into the lower quartile (low expression), interquartile (moderate expression), and upper quartile (high expression) on the log scale.

#### Identifying codons enriched in the 5’-end

To identify codons enriched in the 5’-end and 3’-end, we used both an absolute and relative definition of the termini. The absolute definition considered the termini to be the first and last 100 codons of a coding sequence. In this case, we restricted our analysis to coding sequences with a minimum of 250 codons. The relative definition considered the termini to be the first and last 25% of codons along a coding sequence. In this case, no minimum length cutoff was used.

To identify codons enriched in the 5’-end, an empirical expectation was determined by calculating the frequency of each codon, relative to its synonyms, in the “middle” (i.e., neither of the termini) of the coding sequences across all genes. For each codon, we used a one-sided binomial test to determine if the observed frequency in the 5’-ends was greater than expected by chance based on its observed frequency in the middle of the coding sequence. A similar enrichment test was used for the 3’-ends.

### Calculating the cost of nonsense errors

To better understand the potential importance of nonsense errors in mRNA translation, we calculated the expected cost of producing a complete, functional protein (in terms of the number of hydrolyzed NTPs) E[NTP/protein] from a given transcript that includes the impact of nonsense errors. For simplicity, we exclude the cost of amino acid synthesis from our calculations.

Conceptually, we break the cost of protein synthesis into two overlapping sets of categories: direct vs. indirect costs and fixed vs. variable costs. Direct costs include the NTP used in assembling the small and large ribosome subunits on the mRNA and each elongation step by the ribosome, whereas indirect costs are based on the synthesis cost of the mRNA translation infrastructure. In terms of fixed and variable costs, fixed costs are the direct and indirect costs of translating a protein in the absence of a nonsense error, i.e., costs incurred every round of successful translation. Variable costs are the expected direct and indirect costs of mRNA translation that are wasted when a nonsense error occurs – these costs are variable because they are only incurred if a nonsense error occurs. As a result, the total cost of producing a particular protein E[NTP/protein] is the sum of fixed and variable costs, each of which is comprised of direct and indirect costs.

From our model, it follows that

Fixed Cost = Cost of synthesizing a complete protein

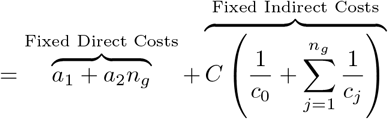

Variable Cost = Expected Number of NSEs/Complete protein *×* Expected cost of a NSE

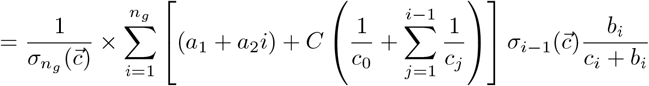

Total Cost = Fixed Cost + Variable Cost

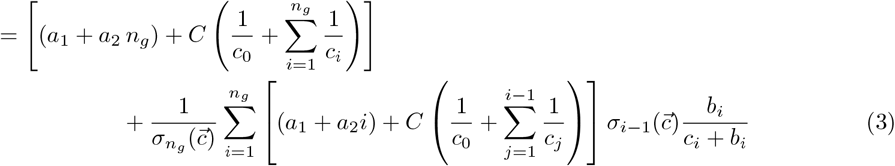

Where *a*_1_ = 4NTP and *a*_2_ = 4NTP are the direct cost of translation initiation and elongation, respectively in terms of hydrolyzed phosphate bonds [18].

Combining the contribution of direct costs to the fixed and variable costs of protein synthesis yields,

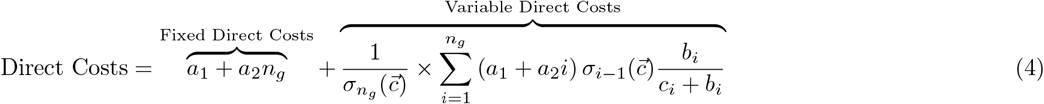

Similarly, combining the contribution of indirect costs to the fixed and variable costs of protein synthesis yields,

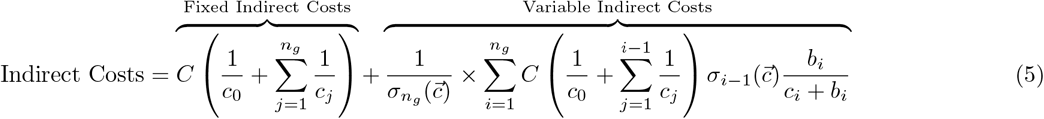

The term 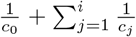 represents the <u>indirect</u> cost of ribosome pausing up to codon *i*. Because RPF data lacks information on the absolute rate of ribosome elongation, we rescaled our estimates of pausing times such that the average elongation rate across the 61 sense codons was ∼ 9.3 codon/sec or, equivalently, 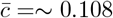 sec/codon. Assuming the average mRNA has a coding sequence of 400 codons, the same average elongation rate as above, and that at any given time 80% of the ribosome population is engaged in mRNA translation [27], it follows that the expected initiation rate is *c*_0_ is ∼ 0.0926/sec (see Supplementary Material for more details).

The parameter *C* converts the indirect cost of ribosome pausing (in units of seconds) into their equivalent costs in terms of NTP and has the units of NTP/sec. Although we are unaware of any empirical estimates of *C*, we use two different approaches to estimate *C* that vary by less than one order of magnitude. One estimate of *C* is based on selection coefficients estimated from a ROC-SEMPPR analysis of the yeast genome and yields *C*_ROC_ = 0.76 NTPs/sec). The second estimate of *C* is based on the assembly cost (in NTPs) and average lifespan (in seconds) of the ribosome and yields 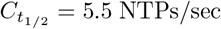. See Supplementary Materials for detailed descriptions of these calculations. Given these two independent estimates of *C* are the only ones we have, we will treat *C*_ROC_ and 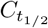 as lower and upper bounds on *C*, respectively. Thus the average cost of ribosome pausing is approximately 0.108sec/codon *× C* = 0.082 or 0.59NTP/codon for *C*_ROC_ and 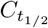, respectively, suggesting indirect costs are a fraction of the direct cost of 4 NTP/codon.

In addition to using our cost estimates directly, we also used them to test the hypothesis that synonymous codon order along a transcript shows evidence of adaptation to reduce the cost of nonsense errors. To do so, we generated a null distribution of the expected cost of mRNA translation for each gene when there is no adaptation to reduce the cost of nonsense errors by permuting the order of its synonymous codon within the coding sequence. These permuted sequences had the exact same amino acid usage, ordering, and codon usage as the sequence observed in the genome, and only differed amongst the ordering of synonyms within each amino acid. For each permutation, we calculated its expected protein production cost using Eq (3). We then compared the mean protein production cost of our population of permutated sequences to that of the protein production cost of the observed sequence. We then scored each observed sequence as having a protein production cost either greater than (0) or less than (1) the expected cost under the null hypothesis. To test whether the observed sequences are biased towards lower costs than expected under the null hypothesis, we performed a binomial test (*H*_0_ : *p* = 0.5). This allowed us to test if the number of sequences with an expected cost less than the mean expected cost across the permuted sequences was greater than expected by chance. Because the gross cost of nonsense errors scales with the protein production rate of a gene, we performed a logistic regression to test if the probability the observed gene has a lower cost than expected increases with gene expression.

### Analysis of other yeasts ribosome profiling datasets

To complement our analysis of the Weinberg et al. data, which is generally considered to be high quality, we also analyzed data from two more recent ribosome profiling experiments: Wu et al. 2019 (SRA: SRR7241903) [29] and Ferguson et al. 2023 (SRA:SRR23242245, SRR23242246) [51] for comparison. These two datasets were processed using the riboviz2 pipeline, with minor differences in the read lengths selected and in defining the A-site (see Supplemental Materials and Methods).

## Results

After PANSE was fit to *S. cerevisiae* ribosome profiling data from Weinberg et al. [27], we compared parameter estimates to appropriate empirical and theoretical proxies, finding them in good agreement (Fig. 1). Both mRNA abundances and tRNA gene copy number (tGCN) are common empirical proxies for translation initiation rates and elongation rates, respectively [24, 27]. We expected PANSE estimates of gene-specific translation initiation rates 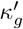 and codon-specific elongation rates *c* to be well-correlated with these empirical proxies. Indeed, translation initiation rates 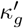 and independent RNA-Seq-based estimates of mRNA abundances are strongly correlated [27] (Fig. 1A, Spearman rank correlation *ρ* = 0.93, *p* < 2.2*E* −16). PANSE estimates of elongation rates *c* and estimates based on tGCNs (including effects of wobble base pairing) are reasonably well-correlated (Fig. 1B, Spearman rank correlation *ρ* = 0.53, *p* = 9.7*E* − 06).

**Figure 1:**
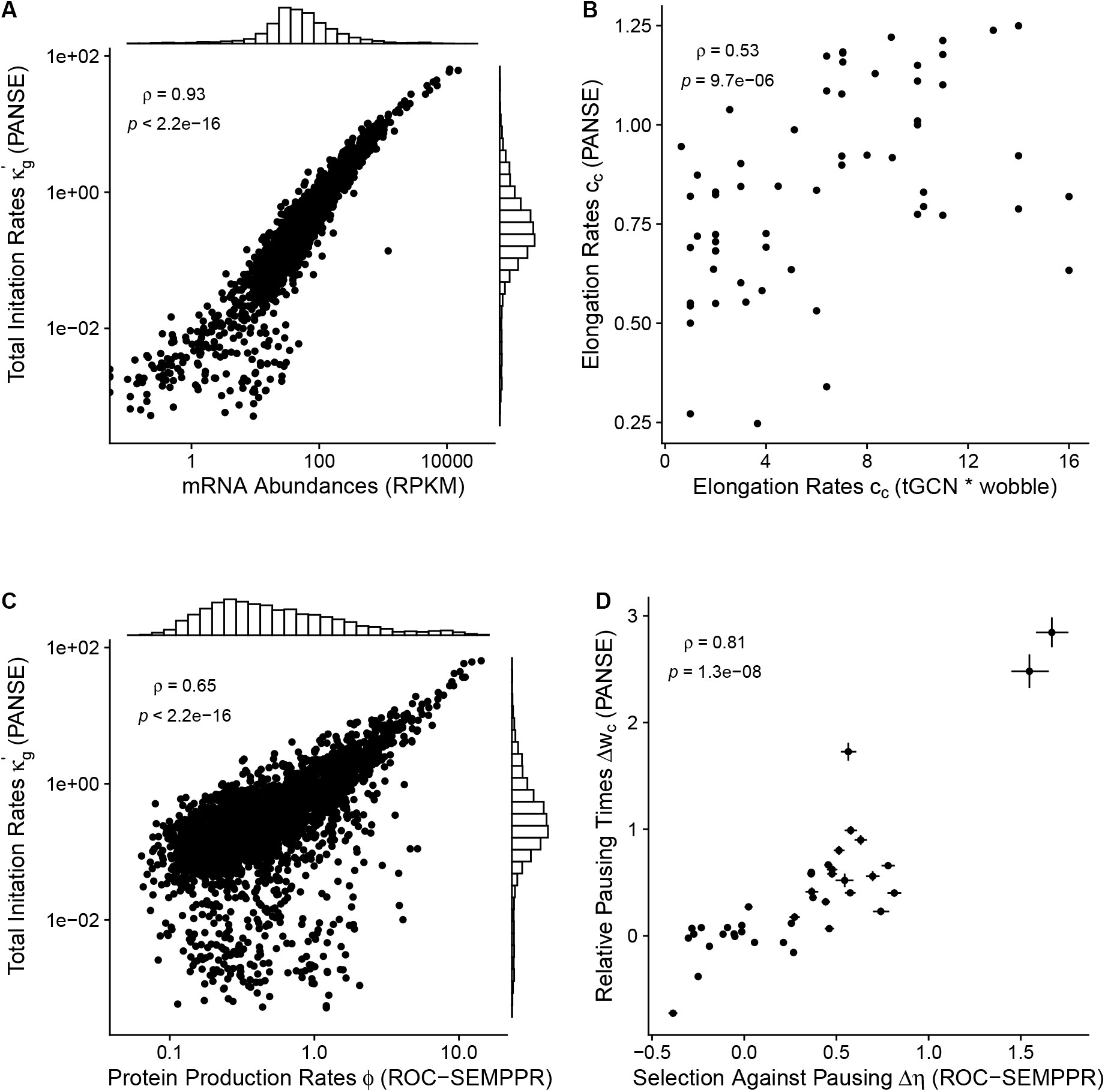
Comparing PANSE gene-specific initiation rates and codon-specific elongation rates to empirical and codon-based estimates. Histograms on x and y-axes represent the distributions of the relevant variables. (A) RNA-seq estimates of mRNA abundances (RPKM) (a common proxy for translation initiation rates) and PANSE estimates of initiation rates 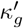. (B) tRNA gene copy numbers (tGCN) based estimates of elongation rates and PANSE estimates of elongation rates. (C) ROC-SEMPPR estimates of protein production rates *ϕ*_*g*_ (based on differences in codon usage) and PANSE estimates of initiation rates 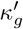. (D) ROC-SEMPPR estimates of selection coefficients Δ*η* and relative ribosome waiting times Δ*w*. Waiting times *w* are defined as the inverse of the elongation rates *w* = 1/*c*. Selection coefficients and waiting times were set relative to the alphabetically last codon for each amino acid.

Theoretical biophysical models rooted in population genetics principles effectively estimate parameters relevant to the evolution of codon usage bias [6, 7, 18, 52]. One such model, ROC-SEMPPR, estimates protein production rates *ϕ* and selection coefficients Δ*η* solely from genome-wide patterns of codon usage bias. ROC-SEMPPR assumes differences in natural selection on codon usage are due to differences in elongation rates *c* between synonyms [6, 7]. We obtained parameter estimates from a previous application of ROC-SEMPPR to the *S. cerevisiae* CDSs [50]. We note protein production rates 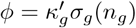, but ROC-SEMPPR ignores translation errors such that *σ*_*g*_(*n*_*g*_) = 1. As expected, initiation rates 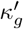 and protein production rates *ϕ* are strongly correlated (Fig. 1C, Spearman rank correlation *ρ* = 0.65, *p* < 2.2*E* − 16). Similarly, estimated elongation rates *c* between synonymous codons and selection coefficient Δ*η* are strongly correlated (Fig. 1D, Spearman rank correlation *ρ* = 0.81, *p* < 1.5*E* − 08). To make elongation rates comparable to the ROC-SEMPPR selection coefficients – which reflect selection against slow codons – we converted our elongation rates to pausing times *w* = 1/*c* and made them relative to ROC-SEMPPR’s pre-defined reference codon for each synonymous codon family, i.e. 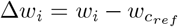, where *i* is a codon and*c*_*ref*_ is the reference synonym.

### NSE rates vary across codons

Theoretical and computational studies often ignore nonsense errors or assume a uniform (background) NSE rate *b* across all codons [21, 30, 33, 34]. However, empirical studies indicate NSEs occur at appreciable frequency [9, 32], with targeted studies suggesting background NSE rates vary across codons [35, 36]. This naturally leads to two questions: do we see evidence of NSEs in ribosome profiling data and (2) do we see evidence that NSE rates (and not just NSE probabilities) vary by codon? To answer question (1) we compared the most complex model (NSE rates vary across codons) to the simplest model (no nonsense errors) using a standard model comparison approach based on the Deviance Information Criterion (DIC) [49, 53]. We found that the variable NSE rate *b* model better fit the ribosome profiling data compared to the no NSE model by 2381 DIC units, indicating the effects of NSEs are detectable through ribosome profiling data and changes to ribosome density along transcripts is not solely due to differences in codon waiting times.

Next, we compared the model allowing NSE rates *b* to vary across codons to a model that assumed all codons had a uniform NSE rate *b*. The uniform NSE rate model still allows for differences in NSE probabilities Pr(NSE) through differences in codon elongation rates. We found that the model allowing for variation in NSE rates *b* across codons is 187 DIC units better than the model assuming uniform NSE rates, indicating strong support for differences in the NSE rates across codons. This suggests the existence of intrinsic factors of codons unrelated to translation speed that alter the probability of NSEs Pr(NSE). NSE rates *b* varied over multiple orders of magnitude, ranging from 8.65 *×* 10^−6^ to 1.98 *×* 10^−3^ (Fig. 2A). When accounting for differences in elongation rates *c*, the mean NSE probabilities Pr(NSE) across codons were on the order of 10^−4^, consistent with previous estimates in *E. coli* and yeast [19–23].

**Figure 2:**
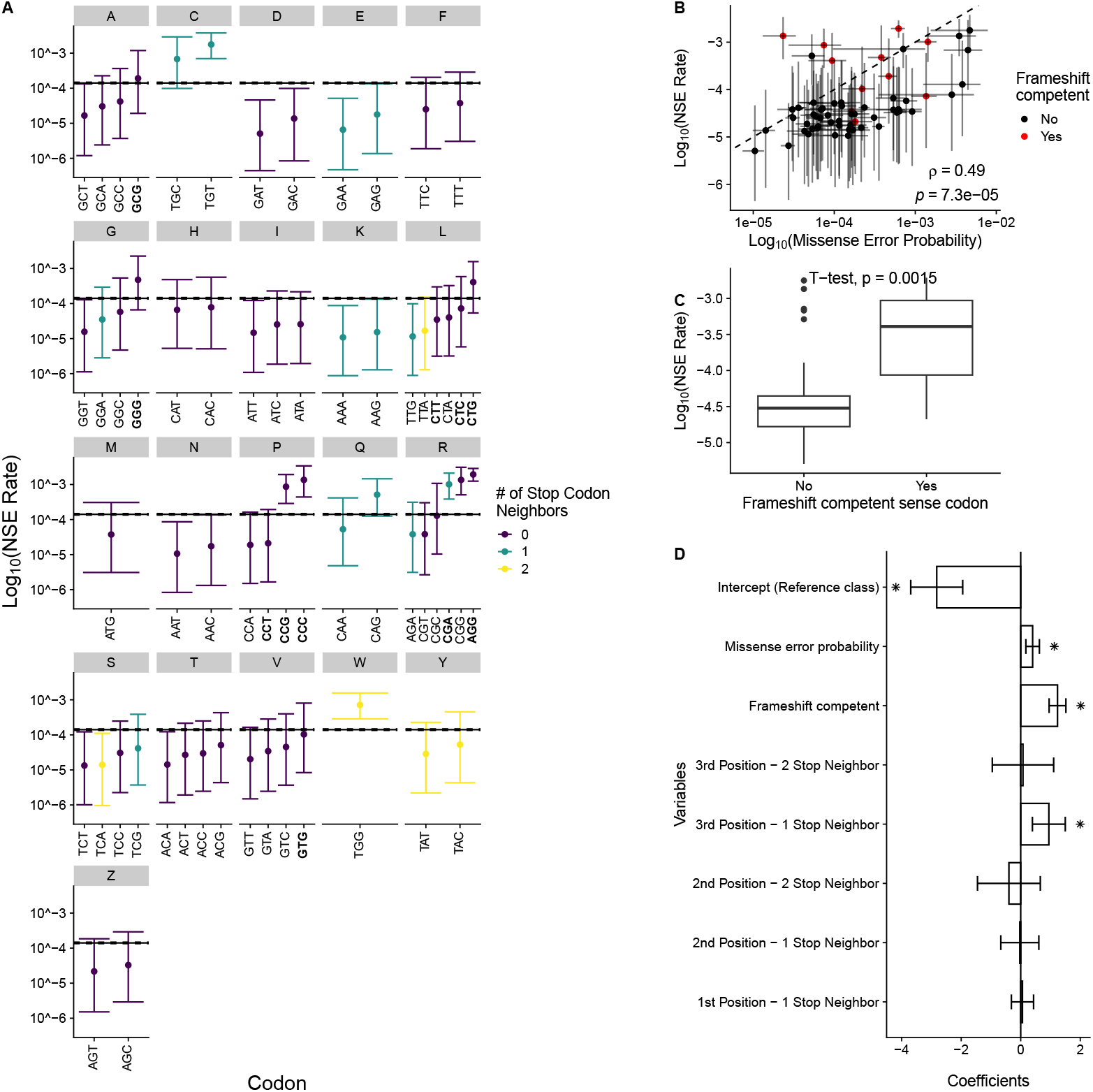
The background NSE Rates *b* vary across codons. (A) Posterior means and 95% highest density intervals (HDIs) of log_10_(NSE Rates *b*) estimated for each codon. Colors indicate the number of nucleotide mutations away a codon is from a stop codon. Solid and dashed black lines indicate the NSE rate *b* posterior mean and 95% HDI for a model fit sharing information across codons (i.e., assuming no variation exists in NSE Rates). To be consistent with our previous work, we separated the amino acid serine into two blocks denoted S and Z. Bolded x-axis labels indicate codons that are frameshift competent [41]. (B) Comparison of NSE rates *b* with empirical missense error probabilities estimated from mass spectrometry data [56]. Spearman rank correlation *ρ* is reported. (C) Comparison NSE rates *b* between 11 frameshift competent codons (as defined in [41]) and the 53 other codons. A Welch’s two sample t-test is reported. (D) Regression coefficient estimates from a weighted multiple regression of codon properties and the NSE rates *b* (on the log10 scale). Variables include the number of stop codon neighbors at each position, whether or not the codon was prone to frameshifts (frameshift competent), and the missense error probability of the codon. The intercept reflects the mean NSE rate *b* of the reference class of codon. Error bars represent 95% confidence intervals. An * indicates statistical significance (*p* < 0.05.).

Importantly, our model is agnostic to the specific mechanisms associated with nonsense errors. Notable mechanisms that can lead to nonsense errors are release factor binding of sense codons, peptidyl-tRNA drop-off, and frameshift errors [54]. Mismatches between the codon and anticodon resulting from near-cognate or even non-cognate tRNA binding (i.e., missense errors) can increase the chances of peptidyl-tRNA drop-off and frameshift errors [39, 42, 55]. Consistent with this, we find that our NSE rate *b* estimates from ribosome profiling are positively-correlated with codon-specific missense error probabilities estimated from a large number of mass spectrometry measurements in *S. cerevisiae* (Spearman rank correlation *ρ* = 0.49, *p* = 7.3*E*−05, Figure 2B) [56]. Previous work in *S. cerevisiae* identified 11 sense codons that were particularly prone to causing frameshifts when located in the P-site of a ribosome, particularly when followed by a slow codon [41]. Based on our model estimates, these 11 “frameshift competent” codons had generally higher NSE rates *b* than the other 53 sense codons (on the log-scale, mean of -3.58 vs. -4.44, Welch Two Sample t-test, *p* = 0.0015, Figure 2C).

To better understand how codon properties associated with higher NSE rates *b* individually contribute to the observed variation, we regressed our estimates against the number of stop codon neighbors (the number of stop codons that are a single nucleotide mutation away from a codon), the average missense error probability, and whether or not the codon is frameshift competent.” (Fig. 2D). We note that we treated the number of stop codon neighbors as a categorical variable, as it is always either 0, 1, or 2. As this is regressing multiple categorical variables against the log_10_(NSE rate *b*), each of these coefficient represents a change relative to a reference class defined by the intercept.

One might expect that having more stop codon neighbors increases the NSE rate *b* of a codon, as there would seem to be a greater chance of being mistakenly recognized by a release factor. Indeed, we find that codons with a single stop codon neighbor at the 3rd position of the codon have an NSE rate *b* almost 1 order of magnitude greater than codons with no stop codon neighbors at the third position (*β* = 0.95, *p* = 0.0012). No such relationship is observed for codons with 2-stop codon neighbors at the third position (*β* = 0.075, *p* = 0.884). In contrast, codons with a stop codon neighbor at either the 1st or 2nd nucleotide position showed no significant difference in NSE rates *b* compared to codons with no stop codon neighbors at these positions. This likely reflects that the ribosome more closely monitors codon-anticodon base pairing at the first 2 positions [57]. Consistent with the independent tests (Figure 2B,C) and in line with the suspected role of missense errors in contributing to peptidyl-tRNA drop-off and frameshifts, we find a positive effect of a codon’s missense error probability (*β* = 0.40, *p* = 8.95*E* − 4) and frameshift competency (*β* = 1.24, *p* = 5.94*E* − 12) on NSE rates *b*.

### Nonsense errors are an unlikely explanation for the “5’-ramp”

Ribosome profiling measurements often exhibit higher ribosome densities at the 5’-end relative to the remainder of the transcript [27]. We simulated ribosome profiling data based on the PANSE model using the parameters estimated from the real data. We observe a moderate correlation between the real and simulated ribosome counts (Spearman rank correlation *ρ* = 0.56, *p* < 2.2*E* − 16). On a position-by-position basis (ignoring the first 200 codons), the average log-fold difference between the real and simulated data was approximately 0.015 (One-sample t-test *p* < 2.2*E* − 16, S4 Fig), suggesting our PANSE model slightly underestimates the number of counts by about 1.5%, on average.

Based on the metagene profile of the ribosome densities for the real and simulated data, we observed good agreement in the post-200^*th*^ codon region (Fig. 3A, unshaded region). This is expected because this was the region used for calculating the likelihood of the data during model fitting. In contrast, the model poorly predicts the ribosome densities in the 5’-region – often called the 5’-ramp region – excluded during the model fitting process. The gradual decrease in ribosome densities along the first 200 codons is far more drastic for the real data than expected based on the model parameters from the remainder of the genes (Fig. 3A, shaded region). This suggests there may be other factors at play in the 5’-ramp that impact ribosome densities, including that this is a technical artifact [27].

**Figure 3:**
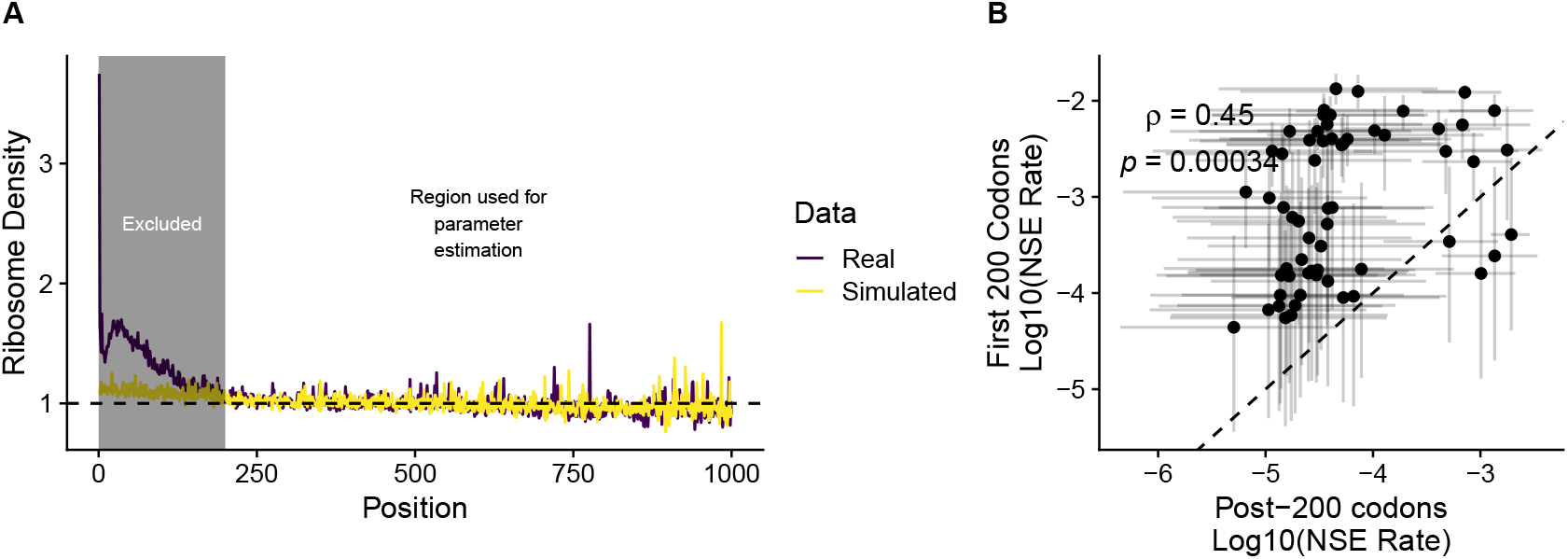
The effects of increased ribosome density at the 5’-end of transcripts. (A) Comparing metagene profiles for real ribosome profiling data (purple) and simulated ribosome profiling data based on the PANSE model (yellow). The first 200 codons are shaded to emphasize these were excluded in the likelihood calculation during model fitting. (B) Comparison of NSE rate estimates for each of the 61 sense codons based on either the first 200 codons or the remainder of the gene.

The discrepancy between real and simulated data raises the question of what NSE rates *b* would be necessary to generate the dramatic drop in ribosome density at the 5’-end. We fit PANSE to only the first 200 codons to determine if the parameters estimated were biologically plausible. Although initiation rates 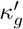 and elongation rates *c* are in good agreement with estimates from the post-200 codon fits (S5 Fig), the NSE rates *b* are significantly greater, with a mean NSE rate of approximately 0.003 across codons (Fig. 3B). This higher NSE rate *b* translates to a higher mean NSE probability Pr(NSE) of approximately 0.004 across codons. This would mean the probability of an NSE occurring within the first 200 codons is approximately 55%, which is unrealistically high. Taken together, our results suggest the 5’-ramp in ribosome profiling data is, at best, only partially due to nonsense errors.

### The probability that translation is completed varies greatly between transcripts

Based on our best model fit, codons differ in their elongation rates *c* and NSE rates *b*, meaning they also differ in their NSE probability (i.e., Pr(NSE) = *b/*(*b*+*c*)) As such, variation in codon usage across CDSs is expected to lead to variation in the probability of a ribosome completing translation *σ*_*g*_(*n*_*g*_). Across all genes, the median probability of a ribosome completing translation *σ*_*g*_(*n*_*g*_) is approximately 0.92 (interquartile range 0.87 – 0.95). A key factor determining the probability of a ribosome completing translation is the length of a transcript. Even if the probability of an NSE at any given codon is rare, longer transcripts afford more opportunities for an NSE to occur. The length of a transcript and its probability of experiencing a nonsense error 1 − *σ*_*g*_(*n*_*g*_) are highly correlated, as expected (Fig. 4A, Spearman rank correlation *ρ* = 0.91, *p* < 2.2*E* − 16). Unsurprisingly, the *S. cerevisiae* proteins YHR099W, YKR054C, and YLR106C are the only transcripts considered with a greater than 50% chance of a ribosome experiencing a nonsense error. This likely has little to do with codon usage, as their respective transcripts have > 3700 codons, such that there are many opportunities for any given ribosome to drop off during elongation.

**Figure 4:**
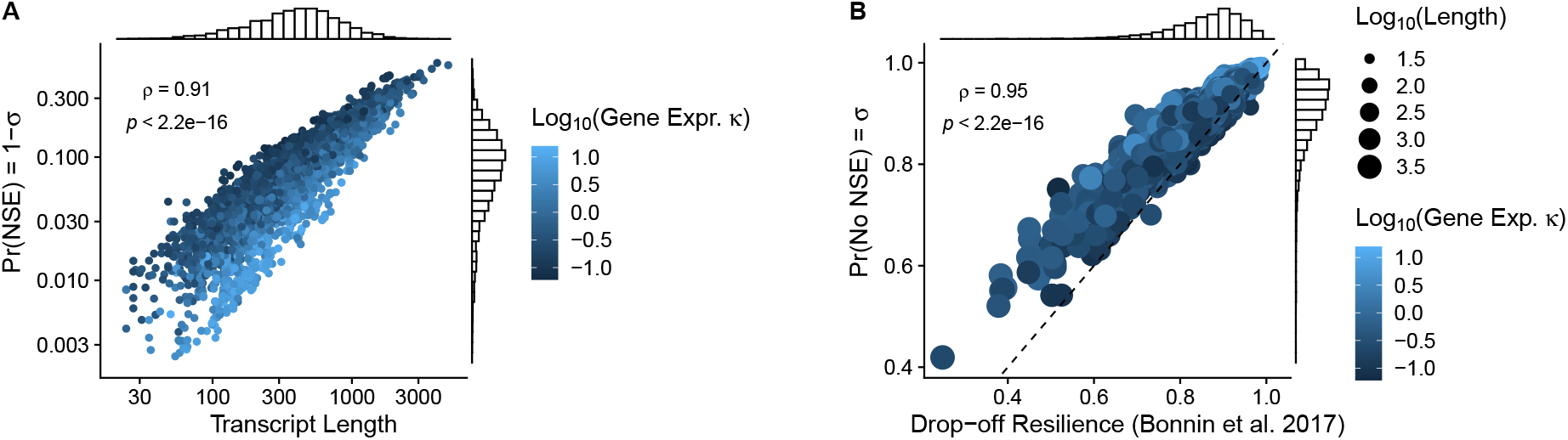
Variation in the probability of a single ribosome completing translation *σ*_*g*_(*n*_*g*_). (A) Relationship between length and the probability of nonsense error occurring, Pr(NSE) = 1 − *σ*_*g*_(*n*_*g*_), with colors representing the ROC-SEMPPR protein production rate *ϕ* for each transcript. (B) Comparison of the probability of a nonsense error not occurring Pr(No NSE) = *σ*_*g*_(*n*_*g*_) with simulation-based estimates of drop-off resilience [34]. Colors represent *κ*_*g*_ from ROC-SEMPPR (i.e. *ϕ*) for each transcript and the size of the point represents the transcript’s length.

A previous study estimated the probability a ribosome completes translation using a Totally Asymmetric Exclusion Process (TASEP) simulation parameterized by polysome profiling data (translation initiation rates), tRNA counts (codon-specific elongation rates), and a single NSE rate estimate from *E. coli* [34]. We compared our estimates of translation completion probability based on a model fit to empirical data *σ*_*g*_(*n*_*g*_) to theoretical expectations based on this TASEP simulation (referred to as “drop-off resilience”), finding them to be highly correlated (Fig. 4B, Spearman rank correlation *ρ* = 0.95, *p* < 2.2*E* − 16). We still observed a high correlation between the empirical and theoretical estimates when conditioning on transcript length (partial Spearman rank correlation, *ρ*_*partial*_ = 0.74, *p* < 2.2*E* − 16), indicating transcript length is not the only cause of the similarity between the two estimates of translation completion probabilities. Estimates of *σ*_*g*_(*n*_*g*_) from PANSE are generally greater than expected based on these TASEP simulations, which assumed a uniform NSE rate for all codons. The NSE rate used for the TASEP simulations was taken from a study that implicitly assumed the drop in ribosome density along transcripts was solely due to nonsense errors [21]. As we and others have shown, the 5’-ramp region inflates estimates of NSE rates [22]. Regardless, the high correlation between *σ*_*g*_(*n*_*g*_) and theoretical expectations suggests our PANSE model generally captures the across-transcript variability in the probability a ribosome completes translation based solely on empirical data.

### Evidence supports adaptation to reduce NSEs

As protein synthesis is energetically costly, natural selection is expected to result in genome-level patterns consistent with adaptations to reduce the costs of NSEs. We specifically examine two key adaptations: (1) reduced frequency of codons with higher NSE probabilities Pr(NSE) moving along the transcript in the 5’ to 3’ direction and (2) an anti-correlation between the frequencies of codons with higher NSE probabilities and gene expression.

#### Evidence that adaptation increases with position

As the energetic investment in producing a protein increases as amino acids are added to the peptide chain, natural selection against NSEs is expected to be weakest near the start of mRNA translation, resulting in position-specific patterns of codon usage in which nonsense error-prone codons are biased toward the 5’-end of a CDS[17, 18, 30, 33, 58]. As such, codons with higher NSE probabilities Pr(NSE) are expected to be enriched in the 5’-ends relative to the remainder of the sequences. To control for the different lengths of CDSs, we assigned codons to the 5’-end based on their relative positions (i.e., their actual position divided by the number of codons in a given CDS) using a relative position cutoff of 0.25. The “middle” of CDSs were similarly defined as those falling between 0.25 and 0.75. By comparing synonymous codon frequencies in the 5’-ends of CDSs compared to the middle, we identified codons enriched at the 5’-end (one-sided binomial test, Benjamini-Hochberg adjusted *p* < 0.05). As expected, codons enriched in the 5’-ends had higher Pr(NSE) than codons showing no difference between the 5’-end and the middle of the CDSs(Wilcoxon rank sum test, *p* = 3.7*E* − 08), Fig. 5A). We observed no such pattern if comparing the Pr(NSE) of codons enriched in the 3’-ends to those not enriched relative to the middle of CDSs(Wilcoxon rank sum test *p* = 0.67, Fig. 5B). We obtained a similar result if we defined the termini as the first and last 100 codons of each CDS (S6 Fig). We emphasize that the first 200 codons of each transcript were not considered in the actual parameter estimation (i.e., were not considered in the likelihood calculation).

**Figure 5:**
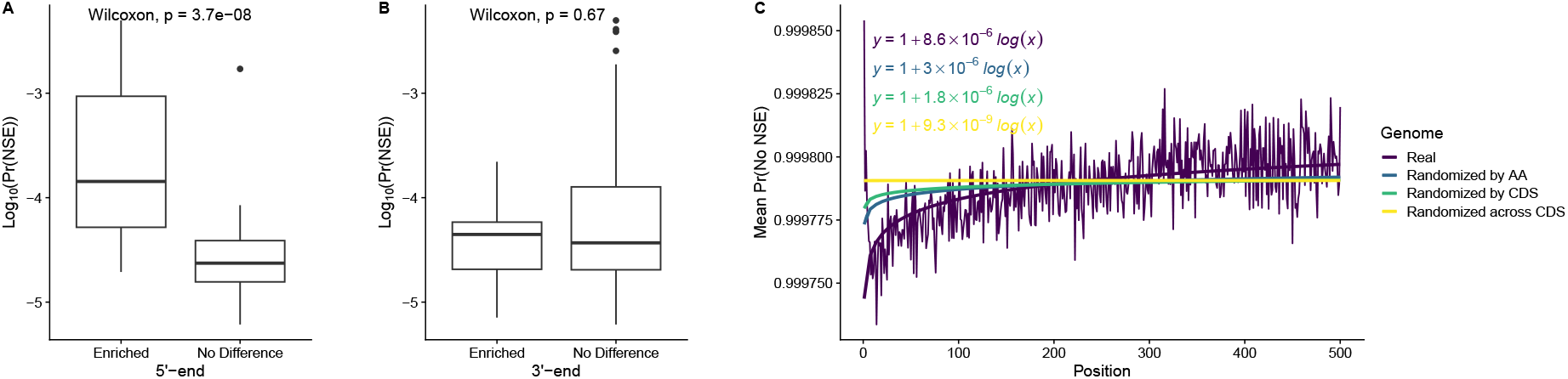
Selection against nonsense errors is weakest at the 5’-ends of transcripts. (A) Comparison of codon-specific Pr(NSE) (on the log scale) of codons enriched in the 5’-end (the first 25% of codons for each sequence) vs. those not significantly enriched relative to the middle of CDSs. Wilcoxon rank sum test p-value reported. (B) Same as in (A), but using the last 25% of codons. (C) Geometric mean of the probability of successful translation by the ribosome up to the 500th codon across all transcripts for the real CDSs (purple). Solid lines and equations represent the linear regressions relating the log(Position) of a codon to the change in probability of elongation (i.e., not an NSE) for the real sequences and the various nulls. For the null regression lines, the mean slope and the mean intercept across the 1000 permuted sequences were used.

Natural selection against nonsense errors is expected to strengthen along transcript, thereby increasing the probability of successful elongation. Except for the first 10 codons, the probability of elongation increases along the transcript before appearing to plateau (Fig. 5B). This increase in the probability of elongation as a function of codon position is consistent with adaptations to reduce the energetic cost of NSEs. Regarding the apparent decrease in the probability of elongation for the first 10 codons, we note that these results are qualitatively consistent with previous findings that observed a decrease in codon bias immediately following the start codon, followed by a gradual increase [58]. The change in the probability of elongation observed in the true sequences is much greater than observed for the null expectations generated by permuting codons (Fig. 5B and S7 Fig). This is consistent with natural selection against NSEs being generally weaker at the 5’-end.

#### Evidence that adaptation against NSEs increases with gene expression

As highly expressed genes generally undergo more rounds of translation (i.e., take up a larger portion of the cell’s energy budget), a lower probability of completing translation in a highly expressed gene will lead to more wasted NTP. Thus, selection against NSEs should increase with gene expression and with highly expressed genes having the greatest probability of completing translation *σ*_*g*_(*n*_*g*_). In our comparisons of sequence length and the probability an NSE occurs, we observed a clear gradient in this relationship: highly-expressed genes tend to have lower probabilities of experiencing a nonsense error compared to transcripts of similar length but lower expression (Fig. 4A). Using mRNA abundances [27] as an independent measure of gene expression, we find a positive correlation between the probability of completing translation *σ*_*g*_(*n*_*g*_) and gene expression (Spearman rank correlation *ρ* = 0.31, *p* < 2.2*E* − 16, Fig. 6A). This analysis does not control for sequence length, indicating that regardless of length, highly expressed genes are more likely to be successfully translated. By controlling for length using a partial correlation, we found that the correlation between gene expression and the probability of completing translation *σ*_*g*_(*n*_*g*_) increased (partial Spearman rank correlation *ρ*_*partial*_ = 0.49, *p* < 2.2*E* −16). As expected, our results are consistent with selection against NSEs being generally stronger in high-expression genes compared to moderate- or low-expression genes of similar length.

**Figure 6:**
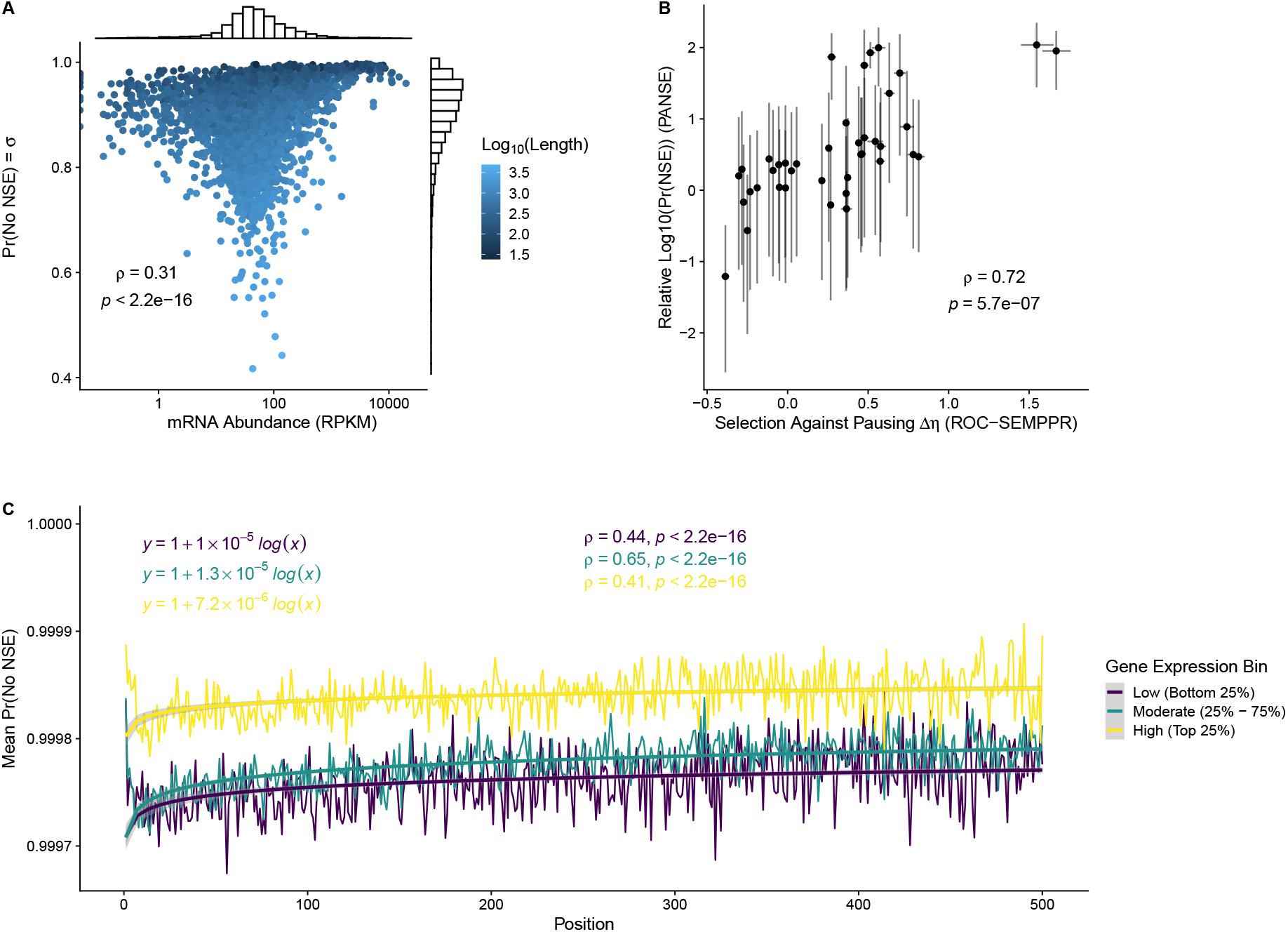
Highly-expressed genes are better adapted to avoid nonsense errors. (A) Comparing gene expression (mRNA abundance RPKM) with the probability of a ribosome completing translation *σ*_*g*_(*n*_*g*_). Histograms on the x and y axes represent the distributions of the corresponding variables. Colors indicate the length of the CDS.(B) Comparing relative NSE rates among synonymous codons to selection coefficients Δ*η* from ROC-SEMPPR. A higher value of Δ*η* indicates a codon that is disfavored by selection relative to a reference codon (the alphabetically last codon for each amino acid). Error bars represent the 95% HDIs. Colors indicate the number of stop codons that can be reached via a single mutation to the codon. (C) Same as in Fig. 5(B) but separating genes into bins based on mRNA abundances (RPKMs). Bins represent the quartiles. Solid lines represent the linear regression relating codon log(position) to the probability of elongation.

ROC-SEMPPR assumes selection for on codon usage is uniform along a CDS, but codons with higher NSE probabilities are expected to be avoided in high-expression genes due to the increased energetic costs of experiencing frequent translation errors. Under selection against NSEs, we expect ROC-SEMPPR’s selection coefficients Δ*η* to be correlated with relative NSE probabilities (i.e., between synonymous codons). We find that Δ*η* and differences in NSE probabilities are well-correlated (Spearman rank correlation *ρ* = 0.72, *p* = 6.7*E* − 07, Fig. 6B), indicating that synonymous codons with higher NSE probabilities are avoided in high-expression genes.

Consistent with the avoidance of synonymous codon with higher NSE probabilities in high-expression genes, we observe the probability of successful elongation by position is greater in high-expression genes (Fig. 6C), indicating these genes are better adapted to reduce the cost of nonsense errors. Independent of gene expression, we observe the probability of successful elongations increases along the transcripts.

### The energetic costs of NSEs are likely substantial

Although NSEs are rarer on average than missense errors [9, 54], they may be more costly due to the high probability that the truncated protein is non-functional. We calculated the expected cost of mRNA translation (in terms of NTP) E[NTP/protein](*c*) (see Materials and Methods) that accounts for the direct ATP cost of translation initiation and peptide elongation, the indirect overhead cost due to ribosome pausing, and the direct and indirect costs of NSEs. The direct and indirect costs associated with translation initiation and elongation are the fixed costs as these will be incurred every time a functional protein is produced. In contrast, direct and indirect costs associated with NSEs as are variable costs as these will only be incurred <u>if</u> an NSE occurs (see Eqn. 3). For the cost of ribosome pausing during translation *C*, we focus on a rough estimate based on ribosome production costs and half-lives 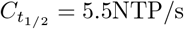 as this estimate is more directly tied to ribosome assembly (see Materials and Methods). Unsurprisingly, the majority of NTP used during mRNA translation is associated with fixed direct costs of translation initiation and ribosome elongation (Fig. 7A); however, this is not expected to impact the evolution of codon usage because fixed direct costs do not vary across codons.

**Figure 7:**
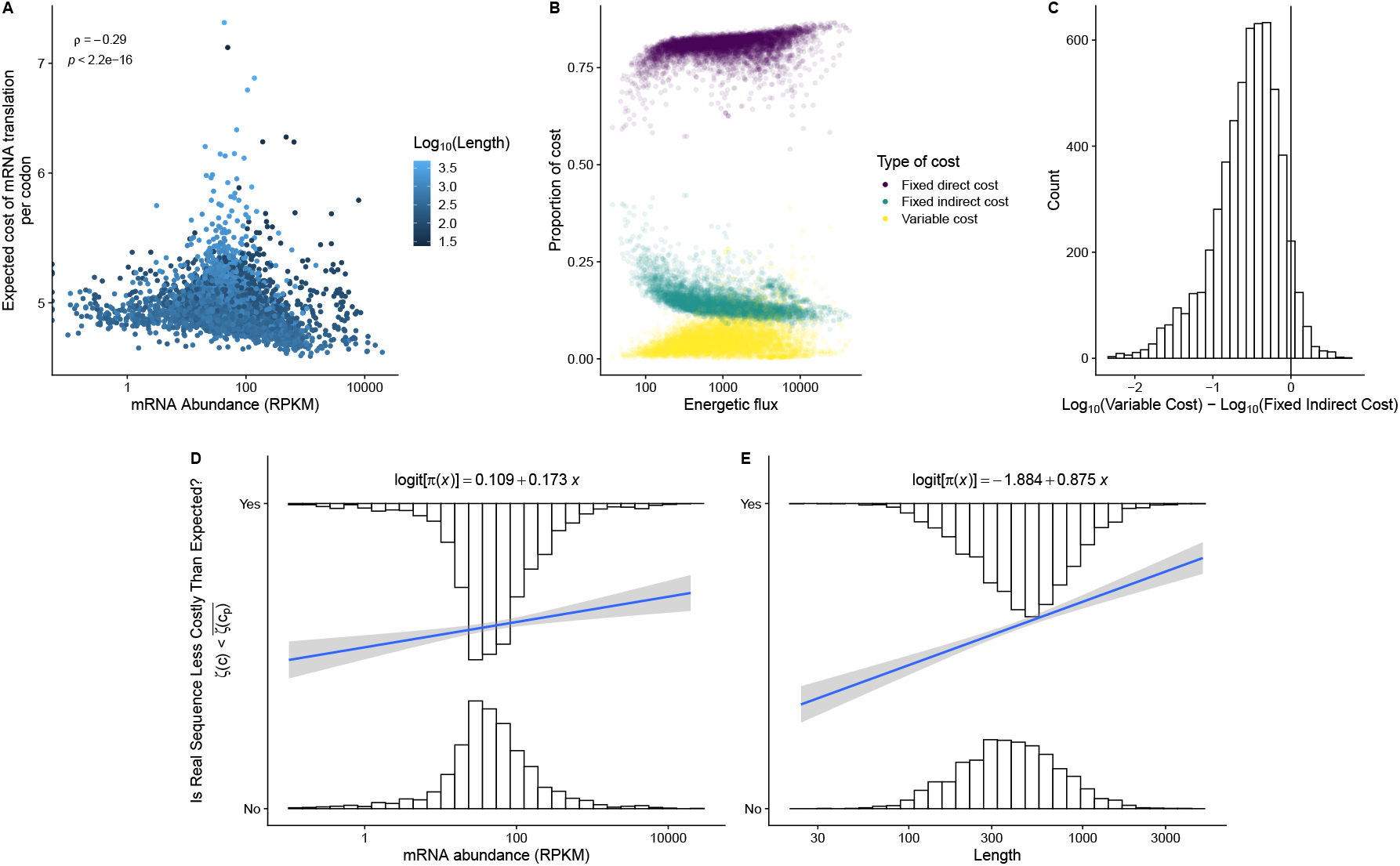
Expected cost of mRNA translation in terms of NTP across genes. (A) Relationship between gene expression and the expected cost of translation per codon. (B) Relative costs of translation in terms of direct costs (translation initiation and peptide elongation), indirect costs (overhead costs of ribosome pausing), and variable costs. Genes are ordered based on total energetic flux: the product of the total costs and protein production rates *ϕ* (as estimated by ROC-SEMPPR). (C) The log fold-difference between the variable costs (i.e. the direct and indirect cost of NSEs) and the fixed indirect cost of elongation (i.e., ribosome pausing). The mean log fold difference is -0.56 (One-sample t-test *p* < 2.2*E* − 16). (D) Relationship between gene expression and the probability of whether or not a gene has a lower observed energetic cost than expected based on the permuted nulls. Histograms represent the distribution of gene expression for genes with and without evidence of adaptation against nonsense errors. The solid line and the equation (slope: represent the result of a logistic regression. The logistic regression slope was statistically significant (*p* = 0.00024), but the intercept was not (*p* = 0.19). (E) Same as in (D), but with length as the predictor variable. Both the logistic regression slope (*p* < 2.2*E* − 16) and intercept (*p* = 2.4*E* − 12) were statistically significant.

Studies on the evolution of codon usage have primarily focused on the cost of pausing, either explicitly or implicitly. Based on 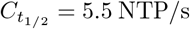, the fixed indirect cost (i.e., the cost of ribosome pausing) is generally greater than variable costs (both variable direct and variable indirect costs) (Fig. 7B). However, variable costs (i.e., the direct and indirect costs associated with an NSE) are usually within an order magnitude of the fixed indirect costs (i.e., the total cost of ribosome pausing for producing a single protein) for the majority of genes under consideration (≈ 86%). We note this is likely a conservative estimate of variable costs. Using *C*_*ROC*_ = 0.7 NTP/s, which is based on comparing ROC-SEMPPR’s selection coefficients Δ*η* to ribosome elongation rates *c*, we found the variable costs to be generally greater than the fixed indirect costs (S8 Fig). Consistent with adaptation to reduce the cost of translation, indirect costs (both fixed and variable) generally decrease as a function of total energetic flux (S9 Fig). We suspect our estimates of *C*_*ROC*_ and 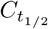 reflect reasonable bounds on the true energetic cost of ribosome pausing. Combined, our results suggest the costs of NSEs are comparable to the cost of ribosomal pausing.

As natural selection to reduce energetic costs is expected to be stronger in high-expression genes, we expect the cost of a gene to decrease as gene expression increases. Consistent with this expectation, we observe the expected cost per codon (i.e. E[NTP/protein](*c*)/*n*, where *n* is the number of codons) is negatively correlated with gene expression (Fig. 7C). To test for evidence of adaptation to reduce the cost of NSEs, we generated a null distribution for the expected translation cost for each CDS based on 1000 permuted sequences of synonymous codons. These permutations could be viewed as nulls reflecting the absence of natural selection against NSEs, as permutations do not alter the codon-order invariant fixed translation costs. As codon usage for *S. cerevisiae* is at selection-mutation-drift equilibrium [6], we are effectively testing against a null that assumes adaptive codon usage via natural selection that varies with gene expression but not with position within a gene e.g., ribosome pausing. We found the true expected cost of a sequence E[NTP/protein](*c*) is less than the mean expected cost of the permuted sequence 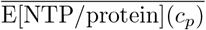 for 59% of CDSs, which is greater than the 50% expected by random chance (two-sided binomial test, *p* < 2.2*E* − 16). The average difference between the true cost and the mean cost of the permuted sequences is approximately 4 NTPs, roughly the same cost as initiating another ribosome or the direct cost of translation elongation.

We have already seen evidence that high-expression genes are better adapted to reduce the cost of NSEs. Furthermore, as NSEs are more likely to occur in longer genes, gene length may be another predictor of adaptation to reduce the cost of NSEs. By classifying CDSs as either those that were better adapted relative to the permuted sequences (i.e., 1 if E[NTP/protein](*c*) 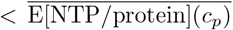 and 0 otherwise), we find highly expressed and longer genes are more likely to be adapted to reduce the cost of NSEs (Fig. 7D,E).

### Parameter estimates across *S. cerevisiae* ribosome profiling datasets are largely consistent

Ribosome profiling, as with any high-throughput experiment, can be subject to technical and biological variation [52]. There are many protocols for performing ribosome profiling, each with their own possible biases [27, 47, 59]. To ensure that our model parameter estimates were generally consistent across measurements, we fit PANSE to data from two independent ribosome profiling measurements: Wu et al. [29] and Ferguson et al. [51]. Although these measurements were performed in *S. cerevisiae*, they used different protocols.

Comparing the metagene plots for the transcripts considered in each of the datasets, Weinberg et al. and Wu et al. exhibit the 5’-ramp; however, this effect is nearly absent from the Ferguson et al. data, with a slight depletion in ribosomes in the region between the 75th and 100 codon (S10 Fig A). We emphasize that none of these reads were included in the actual likelihood estimation. The Ferguson et al. data had the fewest number of transcripts under-consideration (1,918 as compared to 2,785 for Wu et al., and 3,112 for Weinberg et al.). The Ferguson et al. data was also much sparser (S10 Fig B), even if only considering the same sequences across all datasets (S10 Fig C). Despite this, we see overall good agreement between the NSE rate *b* estimates of these two datasets with the Weinberg et al. estimates, despite the reduced number of reads (Spearman rank correlation *ρ* = 0.74 and *ρ* = 0.69 for Wu et al. and Ferguson et al., respectively, S10 Fig D,E).

Notably, the Ferguson et al. dataset has much noisier estimates (S10 Fig E), but this is unsurprising given that it is a much smaller dataset. When assuming the NSE rate *b* is the same across all codons when analyzing Ferguson et al., the NSE rate *b* is lower, but comparable, to the same analysis using the Weinberg et al. data: 9.29 *×* 10^−5^ (95% HDI: 8.27 *×* 10^−5^ − 1.03 *×* 10^−4^) vs. 1.41 *×* 10^−4^ (95% HDI: 1.34 *×* 10^−4^ − 1.47 *×* 10^−4^). Although estimates of ribosome waiting times *w* = 1/*c* (inverse of the elongation rates) were correlated with a proxy based on the tRNA gene copy number, this effect was weakest for the Ferguson et al. data (S10 Fig F). Additionally, the distribution of ribosome waiting times *w* appears narrower than in the Weinberg et al. and Wu et al. data (S10 FigF). Overall, our results suggest we are picking up an overall consistent signal of nonsense errors across these independent datasets, but the reduced number of reads in Ferguson et al. data significantly weakens this signal.

## Discussion

The impact of translation errors on coding-sequence evolution has been a major focus for the last 3 decades, with most of this work focused on the impact of missense errors [10, 12, 16, 60]. Recent advances in mass spectrometry technology and proteome bioinformatics make it possible to detect missense errors on a proteome-scale [13, 56]. Due to the robustness of the genetic code, missense errors may not necessarily lead to a non-functional protein. In contrast, nonsense errors are likely to almost always result in a non-functional protein. There is no current high-throughput technology to directly identify the location of a nonsense error at a particular codon in a transcript. However, high-throughput ribosome profiling allows for a steady-state measurement of the “translatome,” including the ability to assign ribosomes to particular codons [43]. We developed a model of ribosome movement during mRNA translation to quantify codon-specific nonsense error (NSE) rates and probabilities from ribosome profiling data.

Applying our PANSE model to an exemplary measurement for *S. cerevisiae* [27], we found evidence that NSE rates *b* (sometimes referred to as the “background NSE rate”) vary across codons. Prior work generally assumed the probability of a nonsense error occurring at any given codon was solely due to variation in the elongation rates of the codons, with the NSE rate *b* assumed uniform [18, 30, 34]. In contrast, we observed that NSE rates *b* vary over multiple orders of magnitude (10^−6^ to 10^−3^), suggesting other properties of codons contribute to their propensity to trigger a nonsense error aside from differences in elongation rates *c*. Importantly, our model is agnostic to the specific mechanisms that cause nonsense errors; however, our parameter NSE rates *b* reflect three key properties thought to contribute to nonsense error. One potential cause of variation in NSE rates *b* among codons is the propensity for them to be mistakenly bound by release factors [32]. Indeed, codons a single nucleotide away from a stop codon (i.e., has a stop codon neighbor) at the 3rd nucleotide position (cysteine codons TGC/T and tryptophan codons TGG) have a higher NSE rate *b*, on average, compared to codons that do not have a 3rd position stop codon neighbor. The high NSE rate of the tryptophan codon TGG is particularly notable in the context of more direct studies which label this codon a cryptic stop codon due to its ability to bind with eRF1 [36]. Surprisingly, this was not the case for the amino acid tyrosine, for which both of its codons (TAC/T) each have 2 stop codon neighbors (TAA/G) – although the effect of having 2-stop codon neighbors at the 3rd nucleotide is positive compared to a codon with no stop neighbors, it was not statistically significant. Consistent with our results, previous work found eRF1 binds TAC only weakly [36]. Taken together, our results indicate being a single nucleotide away from a stop codon at the wobble position can increase the chances of a nonsense error, but it is not in and of itself sufficient to increase the NSE rates *b*.

We find codons with higher missense error probabilities tend to have higher NSE rates *b*. Previous work suggested mismatches between the codon and anticodon in the P-site can increase the chances of peptidyl-tRNA drop-off and ribosomal frameshifting [38, 39, 42, 55]. Consistent with the latter, codons particularly prone to frameshifts also had higher NSE rates *b*. Recent work in *E. coli* found that peptidyl-tRNA drop-off driven by missense errors generally happens in the subsequent round of elongation [55]. Similar to the incorrect binding of release factors to sense codons [32], slower elongation in the A-site due to e.g., low tRNA availability would seem to increase the probability of peptidyl-tRNA drop-off and ribosomal frameshifting [41].

Other mechanisms may also trigger premature translation termination and be absorbed into the model’s estimates of NSE rates *b* [61–64]. For example, previous work in *E. coli* found that a codon-anticodon mismatch for the tRNA in ribosome P-site was detrimental to the accuracy decoding of the codon in the A-site, increasing the probability of a release factor binding to a sense codon [61, 65] (although we note this was not observed in yeasts using a similar experimental setup [66]). In this case, the probability of a nonsense error at any given codon is not independent, but a function of the missense error rate of its immediate upstream neighbor, among other things. This likely also affects estimates of NSE rates *b* as it pertains to ribosomal frameshifts and peptidyl-tRNA drop-off. The fact that we find codons with higher NSE rates *b* tend to be those with higher missenese error probabilities and higher ribosome frameshift capacities suggests our model detects these effects, but these estimates may be conservative given that we do not consider the surrounding sequence context.

Overall, we observed that simulated ribosome counts across codons and genes generated from PANSE were generally in good agreement with the real data beyond the 5’-ramp region (first 200 codons), indicating PANSE adequately models the underlying processes shaping ribosome profiling data. We emphasize that while the PANSE model does treat elongation rates as a random variable, it does not explicitly account for various potential factors hypothesized to impact local elongation rates, from upstream basic amino acids in the ribosome tunnel to downstream mRNA secondary structure [22, 46, 67]. We note the correlation between the real and simulated data may be inflated due to the latter being generated from parameters estimated from the former, but also attenuated (i.e., biased toward 0) due to the inherent noise in any sequencing data [68]. Regarding the impact of noise, previous work found generally poor to moderate agreement between the ribosome footprint counts at individual codons across independent ribosome profiling experiments [69]. Similarly, for the 3 ribosome profiling experiments we considered, the Spearman rank correlation *ρ* between codon counts on a position-by-position basis ranged from 0.45 (Wu vs. Ferguson) to 0.58 (Weinberg vs. Wu). Future extensions of the model will benefit from explicitly accounting for noise in ribosome profiling data by combining information across independent measurements.

Significant efforts have been made to understand the increased ribosome density at the 5’-end of transcripts observed frequently in ribosome profiling data. In combination with the increased frequency towards slow codons at the 5’-end, previous work hypothesized slow translation was favored at the 5’-end of transcripts to prevent downstream ribosome queuing (“the 5’-ramp hypothesis”) [70]. The 5’-ramp hypothesis remains controversial [27, 71–76]., although recent evidence suggests a very short ramp within the first 5 codons can impact translation efficiency [77]. Our work here is insufficient to directly address the 5’-ramp hypothesis. However, we can say that nonsense errors do not entirely explain the dramatic drop observed in ribosome density at the 5’-end. Given that the expected NSE rates *b* estimated from this region are unrealistically high, we suspect (but cannot confirm) that the increased ribosome density at the 5’-ends results from an experimental artifact [27] and/or a gradual increase in the rate of ribosome elongation over the first 200 codons.

We found numerous lines of evidence consistent with adaptation to reduce the cost of nonsense errors in intragenic codon usage patterns. Generally, selection against nonsense errors is expected to be weakest at the 5’-end of a CDS, as a nonsense error occurring early in translation will waste fewer NTPs. As a result, the codon usage of 5’-ends is expected to be less adapted than regions further along the CDS. Indeed, codons with lower NSE probabilities Pr(NSE) tend to be enriched in the 5’-ends of CDSs. Consistent with this avoidance of higher NSE probability Pr(NSE) codons at the 5’-end, we find that the probability of successful elongation generally increases along a sequence.

In general, high-expression genes tend to avoid codons more prone to nonsense errors. As a result, high-expression genes generally have a higher probability of completing mRNA translation and a lower expected total energetic cost protein production. Furthermore, selection coefficients Δ*η* estimated via the population genetics model ROC-SEMPPR (generally assumed to reflects differences in ribosome pausing times) [7] are well-correlated with relative differences in NSE probabilities Pr(NSE), consistent with high-expression genes avoiding NSE-prone codons. Selection coefficients estimated via ROC-SEMPPR will average over any form of selection pressure that correlates with gene expression [78]. We cannot say to what extent the selection coefficients Δ*η* reflect selection for a reduction in ribosome pausing as opposed to selection against nonsense errors, as both will lead to an increase in translation efficiency.

Surprisingly, moderately expressed genes showed the greatest change along sequences in the probability of a ribosome successfully elongating compared to high and low-expression genes. Assuming selection on codon usage is at least partially due to selection against nonsense errors, this may seem counter-intuitive at first glance. If selection against nonsense errors is strongest in high-expression genes, then selection may be shaping codon usage at the 5’-end. As a result, high-expression genes would be well-adapted even at the 5’-end. In contrast, low-expression genes are expected to be under the weakest selection against nonsense errors, resulting in a lower probability of completion overall that changes very little as a function of position. Moderate-expression genes have may high enough protein production rates such that selection against nonsense errors is expected to be more effective than in low-expression genes, but not so much that the 5’-end is well-adapted. We speculate this explains the greater increase in the probability of elongation along sequences in moderate-expression genes compared to high and low-expression genes.

Although studies on codon usage bias have predominantly focused on selection against ribosome pausing, recent theoretical work concluded the indirect cost of elongation due to ribosome pausing may be less of a factor shaping codon usage bias than other factors, including nonsense errors [67]. Based on a simple model for approximating the energetic costs (in NTPs) for each gene, we found the costs of nonsense errors comparable (i.e., within an order of magnitude) to the fixed indirect costs of ribosome pausing across the majority of the transcriptome. The majority (59%) of genes in *S. cerevisiae* exhibit signals consistent with adaptations to reduce the cost of nonsense errors. However, this is likely a conservative estimate because our permutation test does not represent the expected codon frequencies sans natural selection. More accurately, our test reflects adaptation to reduce the cost of nonsense errors, in which the position of a codon is relevant, to adaptation to reduce a cost invariant with position e.g., ribosome pausing. Although a logistic regression revealed the probability of a gene being adapted increased with gene expression, this was not the case for many high-expression genes (Fig. 7D). As discussed previously, high-expression genes have a relatively low probability of experiencing a nonsense error even at the 5’-end (Fig. 6C), such that permuting synonymous codons within a CDS likely has a small impact on the energetic cost for a significant portion of high-expression genes. Tthe expected energetic cost savings of adaptation against NSEs is approximately 4 NTPs – roughly the same as the cost of initiating a new ribosome for translation or a single round of elongation. Noting that mRNA translation is generally initiation-limited, adaptation against nonsense errors is expected to result in 1 fewer initiation events to produce a functional protein. These estimates were based on many assumptions and parameter estimates from previous model fits, and do not include the potential cost of certain quality control mechanisms that may be triggered by nonsense errors (e.g., nonsense-mediated decay). As such, we suspect our estimates of the energetic cost of nonsense errors are conservative.

We emphasize that our results are consistent with natural selection against nonsense errors, but our parameters do not reflect evolutionary parameters of interest in population genetics studies (e.g., selection coefficients). The most notable pattern of natural selection against nonsense errors is the enrichment of codons with higher NSE probabilities Pr(NSE) in the 5’-end due to presumably weaker selection shortly after translation initiation. As natural selection against nonsense errors is correlated with natural selection for efficient translation, these two selective pressures are correlated across genes, but the latter is not expected to lead to the enrichment of NSE-prone codons in the 5’-end [79].

Other selective pressures are hypothesized to lead to unique patterns of codon usage at the 5’-end of CDSs. A key challenge in understanding selection on synonymous codon usage is the potential for correlated selective effects. First, as previously discussed, is the “5’-ramp hypothesis.” In this case, the enrichment of slow codons, which generally have higher NSE probabilities Pr(NSE), could lead to a similar pattern observed for selection against nonsense errors. However, our previous work using a population genetics model to quantify natural selection shaping codon usage in *S. cerevisiae* and *E. coli* showed that the enrichment in slow codons in the 5’-end is more consistent with a reduction in the strength of natural selection for fast codons, rather than selection generally favoring slow codons in this region [78, 79]. Second, codon usage at the 5’-end is hypothesized to be shaped by selection against mRNA secondary structure to promote efficient translation initiation [71, 80]. However, evidence in *S. cerevisiae* and other species suggests this effect is primarily relevant to the first 10-15 codons [81], whereas we see a gradual decrease in the probability of an NSE over a far broader region in the 5’-end. Third, previous work hypothesized natural selection against ribosomal frameshifting would be stronger at the 5’-end [82]. Ribosomal frameshifts often result in premature translation termination due to encountering an out-of-frame stop codon soon after the frameshift [83–85] and can result from the presence of slow codons in the ribosome A-site in combination with a near- or noncognate codon-anticodon match in the P-site [41, 42, 82]. We expect selection against ribosomal frameshifts to be strongly correlated with selection against nonsense errors. Along these lines, codons prone to causing frameshifts when located in the P-site [41] had higher NSE rates *b*. Future work will focus on quantifying the evolution of codon usage bias as it relates to selection against nonsense errors.

Our results are based on extracting biological information from empirical data using a computational model. As is obligatory, “all models are wrong, but some are useful.” Using the parameter estimate from our model, we were able to test many predictions about expected coding sequence patterns if they were shaped by selection against nonsense errors, using the parameters from our model, many of which were confirmed, illustrating the model’s utility. Our PANSE model of ribosome movement is applicable to any ribosome profiling measurement. This allows for investigations into the impact of different growth conditions on nonsense error probabilities [21, 23]. With the number of species with ribosome profiling measurements growing, it is possible to perform comparative analyses of nonsense error rates and probabilities. Furthermore, using the approaches outlined here, it is possible to quantify the impact of natural selection against nonsense errors on CDS evolution across species and how this may change with e.g., effective population size *N*_*e*_.

## Conclusion

By applying a model of mRNA translation to an exemplary ribosome profiling dataset in *S. cerevisiae*, we found multiple lines of evidence that nonsense errors play a significant role in protein-coding evolution that has largely been underappreciated. Overall, 59% of the CDSs in *S. cerevisiae* exhibit signals of adaptation to reduce the cost of nonsense errors. Natural selection to reduce the cost of ribosome pausing has been the predominant hypothesis to explain codon usage bias, but if the cost of nonsense errors is comparable, if not greater than the cost of pausing, then this hypothesis must be updated or revised. Further consideration of the impact and consequences of nonsense errors is critical for understanding the evolution of codon usage bias, which has been observed to varying degrees across all taxa.

## Data Availability

Raw sequencing reads were obtained from the SRA (SRR1049521, SRR7241903, SRR23242245, SRR23242246). Processed data and scripts for performing model fits and subsequent analyses are available at https://github.com/acope3/Yeast_Nonsense_Error_Analysis.

## Acknowledgements

We thank Premal Shah, Matt Pennell, Edward Wallace, Daohan Jiang, and Josh Schraiber for helpful discussions throughout the course of this project and the writing of this manuscript.

## Supporting information

**S1 Text Supplemental Materials and Methods**.

**S1 Fig Ribogrid from analysis for Weinberg et al. data using the riboviz2 pipeline [48]**. This illustrates the number of ribosome footprints assigned to nucleotide based on the 5’-end of the read. Darker colors indicate more ribosome footprints assigned to a nucleotide. The nucleotide at position 0 indicates the first nucleotide of the start codons.

**S2 Fig Impact of A-site assignment rules on parameter estimates**. Comparison of NSE rate estimates from Weinberg et al. data using either the “standard” A-site 15 nt offsets vs. the offsets estimated by riboWaltz. Spearman rank correlation coefficient *ρ* is reported.

**S3 Fig Factors related to filtering genes from the final analyzed dataset**.(A) Distribution of correlations between position within a gene and ribosome density. (B-C) Genes exhibiting sudden increase in ribosome densities. Dashed lines indicate ATG codons. (D-E) Genes exhibiting a sudden decrease in ribosome densities.

**S4 Fig Deviations between real and simulated ribosome counts across all genes and all positions within the dataset**. Deviations are calculated as the log fold-difference.

**S5 Fig Impact of 5’-ramp region on parameter estimates**. Comparison of (A) elongation waiting times and (B) total initiation rates when considering only the first 200 codons (i.e., the 5’-ramp region) vs. the remainder of the genes. Spearman rank correlations *ρ* are reported. Error bars represent 95% posterior probability intervals.

**S6 Fig First 100 codons are enriched in codons with higher NSE probabilities Pr(NSE)**. Difference in NSE probabilities between codons enriched in the (A) 5’-end and (B) 3’-ends of coding sequences (first and last 100 termini). Wilcoxon rank sum test p-values are reported.

**S7 Fig Null distributions of slope estimates for regression lines relating codon position to the across-gene average in the probability of an NSE per position**. The slope for the real sequences is represented by the dashed line.

**S8 Fig Impact of cost on ribosome pausing** *C* **on total cost estimates**. Comparison of proportion of total cost per gene as a function of total energetic flux (cost times the protein production rate) based on 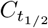 (Assembly) and *C*_ROC_

**S9 Fig Breakdown of energetic costs**. Comparison of proportion of total fixed or variable cost per gene as a function of total energetic flux (cost times the protein production rate) based on *C*_Assembly_ and *C*_ROC_

**S10 Fig Comparison of datasets used as input for PANSE analysis**. (A) Comparison of metagene ribosome densities from Weinberg et al. [27], Wu et al. [29], and Ferguson et al. [51]. (B) Total number of ribosome footprints included in the final PANSE analysis for each dataset (excluding the 5’-ends), expressed as a percentage relative to the Ferguson et al. data. (C) Same as in (B), but only considering the CDSs included in the Ferguson et al. analysis. (D) Comparison of the NSE rate *b* estimates (on log10 scale) from the Weinberg et al. and Wu et al. datasets. Error bars represent the 95% HDIs. (E) Same as in (D), but using the Ferguson et al. dataset. (F) Comparison of PANSE estimated ribosome waiting times *w*_*c*_ and the inverse of codon weights estimated via the tRNA adapation index (tAI).

