## Supplementary material for "The Importance of Nonsense Errors: Estimating the Rate and Implications of Drop-Off Errors during Protein Synthesis": S1 Text

### Supplemental Materials and Methods

#### Model Derivation

Below, we derive the ribosome PAusing and NonSense Error (PANSE) model The goal of our model is to calculate the probability of observing a ribosome profile footprint (RPF) at a given codon in the presence and absence of nonsense errors. The probability of observing a given ribosome footprint is dependent on the population of mRNA polysomes in a cell and calculated under steady state assumptions (i.e. the mRNA and ribosome pools are at steady state and individual mRNAs are at steady state with respect to their polysome distribution).

##### Pausing Time with Nonsense Error Model Definition

We begin our model derivation using the nonsense error (NSE) model of ribosome movement and protein translation derived in (Gilchrist and Wagner, 2006). Our PANSE model differs from this earlier work by (a) allowing the ribosome pausing times for a codon to vary between sites and (b) using a 'waiting time', rather than rate, parameterization.

Starting with eq. 1 in (Gilchrist and Wagner, 2006), we can define the conditional probability of observing an RPF at a given site within an mRNA transcript of a focal gene $g$ as,

$$p_{g,i}|w_{g,i} \propto \kappa_g \sigma(i-1) \frac{w_{g,i} v_{g,i}}{w_{g,i} + v_{g,i}} \tag{1}$$

Where $\kappa_g$ is the translation initiation rate constant for mRNA$_g$ and $v_{g,i}$ and $w_{g,i}$ are the nonsense error and elongation waiting times for codon $i$, respectively. The term $\sigma(i-1)$ describes the probability a ribosome that initiates translation will reach the $i$th codon and is dependent on the $v_{g,j}$ and $w_{g,j}$ for the sites leading up to site $i$ (i.e. $j \leq i-1$). Technically, the translation initiation rate is determined by $\kappa_g \times r$, where $r$ is the density of ribosomes in the cell. However, we assume all mRNAs are equally accessible to ribosomes; as a result, $r$ will cancel out in the following equation and can be ignored throughout. Additionally, we note that $r\kappa_g = J_g$ where $J_g$ is the ribosome flux on an individual mRNA$_g$ per (Pop *et al.*, 2014).

Because the waiting time to an NSE, $v_{g,i}$, is so much greater than the elongation waiting time $w_{g,i}$ we can ignore any actual variation in $v_{g,i}$ between codons of the same type. Thus, while we allow $v_{g,i}$ to be codon specific, we treat each as fixed across positions and genes, using $v_i$ instead of $v_{g,i}$. Further, again because $v_i \gg w_{g,i}$, we can approximate $\frac{w_{g,i} v_i}{w_{g,i}+v_i}$ as $w_{g,i}$ based on a first order Taylor series expansion around $1/v_i = 0$. Taken together, these assumptions allow us to simplify eq. (1) as,

$$p_{g,i}|w_{g,i} \propto \kappa_g \sigma(i-1) w_{g,i}. \tag{2}$$

As mentioned above, the probability a ribosome reaches the $i^{th}$ codon, $\sigma(i-1)$, depends on the probability of successful elongation at the $i-1$ upstream codons. When the waiting time for elongation each position $j$ is known, then

$$\Pr(\text{Ribosome Sampled is at position } j) = \frac{v_j}{w_{g,j} + v_j} \tag{3}$$

and

$$\sigma(i-1) = \prod_{j=1}^{i-1} \frac{v_j}{w_{g,j} + v_j} \tag{4}$$

However, when fitting the model we don't wish to actually estimate individual $w_{g,j}$ values, but instead the parameters of the Gamma $(\alpha_j, \lambda_j)$ distribution that $w_{g,j}$ are drawn from. Thus we treat $w_{g,j}$ as a random variable whose uncertainty we integrate across to get an expected value. Letting $f(\alpha, \lambda)$ represent the PDF of the Gamma distribution, the expectation of the codon specific elongation probability is

$$E\left[\Pr(\text{Ribosome successfully elongates codon } j)\right] = \int_0^\infty \frac{v_j}{w_{g,j} + v_j} f\left(w_{g,j}|\alpha_j, \lambda_j\right) dw_{g,j} \tag{5}$$

$$= \lambda_j v_j \exp\left[\lambda_j v_j\right] E_{p=\alpha_j}\left(\lambda_j v_j\right) \tag{6}$$

where $E_p(z)$ is the generalized exponential integral function (also referred to as Schlomilch functions (Oldham *et al.*, 2009, p.380) and is represented as,

$$E_p(z) = \int_1^\infty \frac{e^{-zt}}{t^p} = z^{p-1} \int_z^\infty \frac{e^{-t}}{t^p} dt = z^{p-1}\Gamma(1-p,z) \tag{7}$$

where $\Gamma(1-p,z)$ is the upper incomplete gamma function (equation 2 in the main text).

Note that, although the complete Gamma function is discontinuous at negative integer values, the upper incomplete Gamma used above is well defined even at negative integers so long as $z > 0$.

Using the above formulations,

$$E\left[\text{Pr(Ribosome successfully elongates codon } j)\right] = \exp\left[\lambda_j v_j\right] \left(\lambda_j v_j\right)^{\alpha_j} \Gamma(1-\alpha_j, \lambda_j v_j) \tag{8}$$

Assuming independence in $w_{g,i}$ between positions means that the expected value of $\sigma_g(i)$ is,

$$\mathrm{E}\left[\sigma(i)\right] = \exp\left[\sum_{j=1}^{i} \lambda_j v_j\right] \prod_{j=1}^{i} \left(\lambda_j v_j E_{\alpha_j}\left(\lambda_j v_j\right)\right) \tag{9}$$

Note that these $\lambda_j$ terms are equivalent to $\lambda_c$.[1] To approximate $\mathrm{E}\left[\sigma_g(i)\right]$, we set $v_i = \bar{v}/k_i$ where $\bar{v}$ is the mean nonsense error waiting time across the set of all sense codons and $k_i$ is a codon $i$ specific scaling factor with $k_i > 0$ and the constraint $\sum_i^n k_i = 1$.

Taking a first-order Taylor Series approximation around $1/\bar{v} = 0$.

$$\mathrm{E}\left[\sigma(i)\right] = 1 - \sum_{j=1}^{i} \frac{\alpha_j}{\lambda_j v_j} + O(1/\bar{v}^2) \tag{10}$$

For completeness, we note that the second-order approximation around $1/\bar{v} = 0$ is,

$$\mathrm{E}\left[\sigma(i)\right] = 1 + \sum_{j=1}^{i} \left[\frac{\alpha_j}{\lambda_j v_j}\left(-1 + \sum_{k=1}^{i} \frac{\alpha_k}{\lambda_k v_k}\right) + \frac{\alpha_j}{{\lambda_j}^2 v_j^2}\right] + O(\bar{v}^3) \tag{11}$$

which can be written in terms of the moments of $w_i$

$$= 1 + \sum_{j=1}^{i} \left[\frac{E\left[w_j\right]}{v_j}\left(-1 + \sum_{k=1}^{i} \frac{E\left[w_k\right]}{v_k}\right) + \frac{\mathrm{Var}\left(w_j\right)}{v_j^2}\right] + O(\bar{v}^3) \tag{12}$$

Note also that there should be a way to represent and, hopefully, efficiently calculate the above equation using matrix notation.

An alternative is to approximate $\ln(\sigma_i)$ around $\bar{v}$ and then exponentiate this approximation. The first-order approximation is,

$$\ln(\mathrm{E}\left[\sigma(i)\right]) = -\sum_{j=1}^{i} \frac{\alpha_j}{\lambda_j v_j} + O(\bar{v}^2) \tag{13}$$

The second-order approximation is,

$$\ln(\mathrm{E}\left[\sigma(i)\right]) = \sum_{j=1}^{i} \left[-\frac{\alpha_j}{\lambda_j v_j} + \frac{\alpha_j}{{\lambda_j}^2 v_j^2} + \frac{1}{2}\left(\frac{\alpha_j}{\lambda_j v_j}\right)^2\right] + O(\bar{v}^3) \tag{14}$$

Assuming independence in sampling, the total probability of a randomly selected footprint is from position $i$, $P_{g,i}$, is

$$P_{g,i} = p_{g,i} m_g / Z \tag{15}$$

[1] $\lambda_c$ is different, but proportional to the term $\lambda'_c$, which we introduce below

where $m_g$ is the density of mRNA$_g$ in the cell and $Z$ is a partition function that ensures our sampling probabilities across the transcriptome sums to 1 and is defined as,

$$Z = \sum_g m_g \left( \sum_i p_{g,i} \right). \tag{16}$$

Equations (15) and (16) indicate that our choices of time and volume units for $\kappa_g$ and $m_g$ on their own are irrelevant, what is relevant is their values relative to one another. This is because our choice of time units for $\kappa_g$ and density for $m_g$ will rescale both $p_{g,i}$ and, thus $Y_{g,i}$ and $Z$. As a result, we choose our time units such that the expected waiting time for methionine is 1, i.e. $E(w_{\text{ATG}}) = 1$.

Although our partition function $Z$ is technically a random variate (and one we estimate for a given dataset), we treat it as a fixed value based on its expected value $Z$ where,

$$E(Z) = E(\sum_g m_g \sum_i p_{g,i}) = \sum_g \kappa_g m_g \sum_c n_{c,g} (\frac{\alpha_c}{\lambda_c}) \tag{17}$$

Letting $Y = \sum_{g,i} Y_{g,i}$ be the footprint sample size, where $Y \gg 1$ and $P_{g,i} \ll 1$, then we can approximate the probability of observing $Y_{g,i}$ samples of a footprint in mRNA$_g$ at position $i$ using a Poisson distribution with a sampling rate of $YP_{g,i}$. Note that deep sequencing may violate the sampling with replacement assumption of the Poisson distribution. Given our assumption about the distribution of $p_{g,i}$, $Y_{g,i} \sim \text{NB}(x = \alpha_c, p = \kappa_g m_g / (\lambda'_c + \kappa_g m_g)$ where $\lambda'_c = \lambda_c Z/Y$.

Taken together, the probability of observing a ribosome footprint at codon position $i$ in gene $g$ is,

$$\Pr\left(Y_{g,i} \middle| \alpha_c, \lambda'_c, \kappa'_g, \text{E}\left[\sigma_g(i-1)\right]\right) = \frac{\Gamma\left(\alpha_c + Y_{g,i}\right)}{\Gamma\left(\alpha_c\right) Y_{g,i}!} \left( \frac{\kappa'_g \, \text{E}\left[\sigma_g(i-1)\right]}{\lambda'_c + \kappa'_g \, \text{E}\left[\sigma_g(i-1)\right]} \right)^{Y_{g,i}} \left( 1 - \frac{\kappa'_g \, \text{E}\left[\sigma_g(i-1)\right]}{\lambda'_c + \kappa'_g \, \text{E}\left[\sigma_g(i-1)\right]} \right)^{\alpha_c} \tag{18}$$

Note, $m_g$ and $\kappa_g$ are gene $g$ and environment specific terms which can be equated to the equilibrium protein synthesis *initiation* rate $\kappa'_g$ for gene $g$ under the experimental conditions, i.e. $\kappa'_g = \kappa_g \; m_g$. The composite parameter $\lambda'_c$ consists of the codon-specific scale term $\lambda_c$ and the ratio of $Z$ to $Y$, two genome-wide parameters. The ratio $Y/Z$ represents as the sampling efficiency of an experiment, the proportion of the RPF state space is sampled. If $Y/Z \ll 1$, then sampling is sparse.

Applying eq. 18 to each codon site within a gene allows us to define a likelihood function L for the focal gene $g$ given the observed footprints from that gene $Y_g$ as,

$$\text{L}\left(\alpha_c, \lambda'_c, \kappa'_g \middle| Y_g\right) = \prod_{i=1}^{n_c} \frac{\Gamma\left(\alpha_c + Y_{g,i}\right)}{\Gamma\left(\alpha_c\right)} \left( \frac{\kappa'_g \, \text{E}\left[\sigma_g(i-1)\right]}{\lambda'_c + \kappa'_g \, \text{E}\left[\sigma_g(i-1)\right]} \right)^{Y_{g,i}} \left( 1 - \frac{\kappa'_g \, \text{E}\left[\sigma_g(i-1)\right]}{\lambda'_c + \kappa'_g \, \text{E}\left[\sigma_g(i-1)\right]} \right)^{\alpha_c} \tag{19}$$

where $n_c$ is the number of codons in the ORF of gene $g$ (equation (1) in the main text). Note we dropped the $Y_g^c!$ term because that will be the same for all parameter values. The total Likelihood of the data is simply

$$\text{L}\left(\alpha_c, \lambda'_c, \kappa'_g \middle| Y\right) = \prod_{g=1}^{n_g} \text{L}\left(\alpha_c, \lambda'_c, \kappa'_g \middle| Y_g\right)$$

Note that $Y_{g,i}!$ has be dropped from our Likelihood equation and the $p_{g,i}$ terms used to calculate $Z$ include $\sigma(i-1)$ terms. Further, note that $Z$ implicitly has the average value of $\kappa'_g$ in it such that our choice of scale for $\kappa'_g$ will, in turn, scale, $Z$. Thus, in this model we have both $\lambda_c$ and $\lambda'_c$ and, as a result, we have an additional, genome-wide parameter to estimate $U = Z/Y$.

For completeness, we define $\phi_g$ as the target production rate of the functionality produced by a complete, error-free protein, i.e. $\phi_g = \kappa'_g \sigma_g(n_g)$ as in Gilchrist (2007). We also note that the rate based formulation of our model with $w_{g,i} = 1/c_{g,i}$, where $c_{g,i}$ is the elongation rate of codon $g, i$, $c_{g,i} \sim \text{InverseGamma}(\alpha_i, \lambda_i)$, and $v_{g,i} = 1/b_{g,i}$ where $b_{g,i}$ is the nonsense error rate of codon $g, i$ results in the exact same likelihood functions as and, hence, . For continuity with our previous work and to aid the interpretation of our results in the main text we present our results using the parameters $c$ and $b$ in the rate formulation of the model.

#### Analysis of more recent ribosome profiling datasets

To complement our primary analysis of the ribosome profiling experiment from Weinberg et al. (Weinberg *et al.*, 2016), we also analyzed ribosome profiling measurments taken from Wu et al. (Wu *et al.*, 2019) and Ferguson et al. (Ferguson *et al.*, 2023). For Wu et al., we used a single replicate that was measured using cycloheximide (CHX) to stall elongation. Characteristic of many of datasets relying on (CHX), we observe a bimodel RFP read length distribution with peaks at 21 and 29 nucleotides (nt), with the latter being much larger. Previous work suggests these different read length reflect ribosomes in different stages of elongation (Lareau *et al.*, 2014; Wu *et al.*, 2019). Unlike with the Weinberg et al. data, for which no bimodal distribution is observed, we included reads of lengths 20-22 in addition to 28-30 nts. Because of this, we also chose to use riboWaltz2 to predict the best A-site offset for each read length (in the form of read length:offset): 20nt:15, 21nt:15, 22nt:16, 28nt:15, 29nt:16, and 30nt:16.

For Ferguson et al., we combined the results from a riboviz2 analyses of the two technical replicates for a ribosome profiling procedure that relied on ordered two-template relay (OTTR) for library generation and P1 as a nuclease (Ferguson *et al.*, 2023). Using this procedure, read lengths are much more broadly distributed, with a peak in the mid-30s (Ferguson *et al.*, 2023). As with Wu et al., we elected to use riboWaltz to predict the best A-site offsets: 30nt:15, 31nt:15, 32nt:15, 33nt:15, 34nt:16, 35nt:16, 36nt:18, and 37nt:19.

##### Filtering of processed ribosome profiling data

After considering only genes that had mapped reads, we made many *post-hoc* filtering decisions to eliminate potential biases or noise that could impact parameter estimation. Genes were removed from the final dataset used for analysis via PANSE if they met any of the following criteria (see also Figure **??**).

1. Showing detectable homology at the nucleotide level based on an all-vs-all BLAST of the yeast transcriptome using default settings.

2. Contain a codon with an abnormally high ribosome density relative to the rest of the transcript, which could indicate ribosome stalls. For each gene, this was determined using a Z-score calculated for each codon based on the log of the ribosome densities within that gene. Any gene that contained a codon with $Z > 3.92$ was excluded from the final dataset.

3. Were in the top 5% (i.e. most positive) of Spearman rank correlations between codon position and ribosome count. Based on manual inspection, many of these genes appeared to have ribosomes initiating at a different start codon than the one annotated, indicating a possible annotation error.

4. Had a greater number of codons with no observed ribosome footprints in the first 10% of the transcript than expected based by chance based on the number of missing codons in the last 25% of the transcript (i.e., a one-sided binomial test with the null determined by the number of codons with 0 ribosome footprints in the last 3'-end of the transcript). This was another test attempting to isolate potential start codon misannotations.

5. Were in the bottom 5% (i.e. most negative) of Spearman rank correlations between codon position and ribosome count. Many of these genes appeared to have abnormalities, such as programmed frameshifts.

6. Were less than 225 codons long.

This ultimately led to final datasets with 3,112 genes (approximately 50% of the protein-coding sequences in *S. cerevisiae*) from Weinberg et al., 2,785 sequences from Wu. et al., and 1,918 sequences from Ferguson et al.

#### Breakdown of energetic costs

##### The indirect costs of mRNA translation

Indirect costs are based on the synthesis cost of the protein translation infrastructure, which we base simply on the cost in NTP of synthesizing the proteins and rRNA found in the large and small ribosome subunits (201,523 NTP) divided by its expected lifespan in the cell (36,360 sec). Note that the 201,523 NTP calculation ignores the cost of nonsense errors and, thus our indirect cost in NTP/sec is likely an underestimate. The indirect cost of protein synthesis (201,523 NTPs) divided by the median lifespan of a ribosomal protein (606 min or, equivalently, 36,360 sec which implies that the indirect cost of each ribosome imposes a cost of 201,523 NTP/36,360 sec $\approx$ 5.5 NTP/sec. Assuming the ribosome spends about 10.8 sec in the cytosol between rounds of translation and a cost of 5.5 NTP/sec corresponds to an indirect cost of translation initiation of 50.4 NTP/initiation.

##### Converting units of elongation waiting times

Because RPF data lacks information on the absolute rate of ribosome elongation, our RFP estimates of $\overline{w}$, which are scaled in time units relative to the methionine codon AUG such that $\overline{w}_{\text{AUG}} = 1$. In order to convert our estimates of $\overline{w}$ into time units of sec, we rescaled our estimates of $\overline{w}$ such that the harmonic mean $\overline{w}$ across all sense codons was 9.3 codons/sec (or equivalently 0.108 sec/codon) (Arava *et al.*, 2003; Shah *et al.*, 2013). (We used the harmonic mean instead of the arithmetic mean because $\overline{w}$ are in units of sec and rates are in units of 1/sec.)

##### Estimating average time between initiation events $w_0$

To estimate the average number of seconds a ribosome spends in the cytosol between rounds of mRNA translation (sec/cytosol), we first assumed that the average protein length is 400 amino acids long. Again assuming codons are translated at an average speed of 9.3 codons/sec it follows that on average the ribosome would spend 400codon/(mRNA translation) $\times$ 0.108sec/codon = 43.0sec/(mRNA translation). Assuming ribosomes are actively translating 80% of the time, it follows that each round of translation takes approximately 43.0sec/(mRNA translation) $\times$ (mRNA translation)/(0.8total) = 53.8 sec/total, of this total time, 20% is spent in the cytosol, thus $w_0$ = 53.8sec/total $\times$ 0.2total = 10.8 sec or, equivalently, 9.26/sec.

##### Estimating the energetic cost of ribosome pausing

As an alternative to quantifying the indirect cost of pausing, a reasonable estimate can be obtained by comparing selection coefficients $\Delta\eta$ from ROC-SEMPPR to waiting times $\overline{w}$ estimated from PANSE. ROC-SEMPPR's estimates of selection coefficients $\Delta\eta$ will reflect an average over different selective pressures on codon usage that scale with gene expression (Cope and Gilchrist, 2022), but it is believed to primarily reflects natural selection for efficient translation (Shah and Gilchrist, 2011; Wallace *et al.*, 2013; Gilchrist *et al.*, 2015). Selection coefficients $\Delta\eta_{i,j}$ between synonymous codons $i$ and $j$ are represented by the equation

$$\Delta\eta_{i,j} = 2qN_e(\eta'_i - \eta'_j)$$

where $N_e$ is the effective population size, $q$ is a scaling term representing the proportional decline in fitness per ATP wasted per unit time (scaled such that the average protein production rate across the genome is 1), and $\eta'$ represents the expected cost of translation (assuming no errors). Using the definition of $\eta'$ defined in our previous work (Shah and Gilchrist, 2011; Gilchrist *et al.*, 2015)

$$\begin{aligned}\eta' &= a_1 + \sum_{i=1}^{n_g} a_2 + C\overline{w}_i \\ \implies \Delta\eta_{i,j} &= 2qN_eC(\overline{w}_i - \overline{w}_j) \\ \implies C &= \frac{\Delta\eta}{2qN_e(\overline{w}_i - \overline{w}_j)}\end{aligned}$$

we can estimate $C$ by comparing elongation waiting times $\overline{w}$ and selection coefficients $\Delta\eta$. Estimates of waiting times via PANSE are not in units of time and are relative to the codon ATG to address identifiability issues.

For the effective population size $N_e$, we used an estimate from *S. paradoxus*, a sister species of *S. cerevisae* ($N_e = 1.36 \times 10^7$), as done previously (Gilchrist, 2007; Shah and Gilchrist, 2011). For the other scaling term $q$, we used a previous estimate $q = 4.19 \times 10^{-7}$(Gilchrist, 2007). We calculated $C$ based on our parameter estimates for each amino acid and used the median value of $C$ as our estimate. We note that $C < 0$ in some cases due to discrepancies between $\Delta\eta$ and $\Delta\overline{w}$, which could arise for several reasons (Shah and Gilchrist, 2011). These were excluded when calculating the median.

#### Important considerations when preparing ribosome profiling data

Although translation initiation rates are generally well approximated solely from the number of ribosome reads mapped to a sequence (Ingolia *et al.*, 2009; Oh *et al.*, 2011; Li *et al.*, 2014; Weinberg *et al.*, 2016), quantification of codon-specific parameters necessitates generally reliable assignment of codons to ribosome A-sites. In eukaryotic ribosome profiling data, most ribosome footprints are between 28 and 30 nucleotides long. with a 15 nucleotide offset generally providing adequate assignment of codons to A-site (Ingolia *et al.*, 2009). We generally expect that a 15 nucleotide offset relative to the 5'-end is sufficient, but future work would benefit from systematically investigating the impact of assigning codons to A-sites. Using predictions from the Bioconductor R package riboWaltz for the Weinberg et al. data, we observed little impact of A-site offset on the NSE rate parameter estimates.

The accurate assignment of reads to codons in the A-site is particularly challenging in ribosome profiling for prokaryotes. Due to the ability of prokaryotic ribosomes to inhibit RNase I, prokaryotic ribosome profiling protocols rely on mononuclease (MNase) to digest unprotected mRNA (Mohammad *et al.*, 2019). MNase is less specific at digesting mRNA, resulting in a broader distribution of read lengths (Li *et al.*, 2014; Woolstenhulme *et al.*, 2015; Mohammad *et al.*, 2016, 2019) that make accurately assigning codons to A-sites more challenging. Although assigning codons to A-sites by mapping relative to the 3'-end of reads appears to be the most reliable approach, the relationship between codon usage and ribosome waiting time remains weak (Mohammad *et al.*, 2019). Due to this greater uncertainty, further caution should be taken when interpreting the results of PANSE (or any model) applied to prokaryotic ribosome profiling data.

Early ribosome profiling measurements performed using cycloheximide protocols showed little correlation between ribosome densities and tRNA abundances (or tRNA gene copy numbers); however, more recent measurements using cycloheximide found a correlation between densities and the tRNA pool consistent with the correlations observed from flash-freeze protocols (Wu *et al.*, 2019). Unlike measurements performed using flash-freeze, cycloheximide-based protocols tend to result in a bimodal distribution of read lengths, with one centered around 28 nucleotides and another around 21 nucleotides (Lareau *et al.*, 2014). Although the former peak is generally the more abundant, the ribosome densities estimated from the 21 nucleotide reads were better correlated with tRNA abundances (Wu *et al.*, 2019).

#### Limitations of the model

A key assumption of the PANSE model is ribosome elongation at each codon is independent of the other codons, meaning it does not explicitly account for ribosome queuing. Under the assumption that translation is initiation-limited, ribosomes are generally expected to be spaced out along an mRNA transcript, with queuing events relatively limited (Plotkin and Kudla, 2011). However, there are numerous examples of paused or stalled ribosomes that may occur for a variety of reasons, including but not limited to slow codon pairs, interactions between the amino acid and ribosome tunnel (Duc and Song, 2018), and mRNA secondary structure. As ribosome queues may result from stalled ribosomes, we removed any genes containing codons with abnormally high ribosome density compared to the transcript overall. Stalled ribosomes may also trigger the no-go decay mechanism in which stalled ribosomes are removed and the transcript is degraded (Presnyak *et al.*, 2015). Future work could incorporate disome ribosome profiling to potentially incorporate ribosome queuing (Zhao *et al.*, 2021). Unlike traditional monosome-based ribosome profiling, disome profiling sequences reads covered by queued ribosomes. That said, our estimates of a ribosome completing translation $\sigma$ were in good agreement with theoretical estimates based on a TASEP framework that allows for ribosome queuing (Bonnin *et al.*, 2017). Based on this, we suspect ribosome queuing generally has little impact on the frequency of nonsense errors.

PANSE assumes the waiting times for each codon follow a gamma distribution, allowing for variability in the elongation rates of a codon instead of assuming a constant value across all occurrences of a codon. Although the identity of the codon at the A-site plays a significant role in determining the elongation waiting time of a ribosome, the upstream and downstream context (e.g., amino acids in the ribosome tunnel, upstream mRNA secondary structure) may also shape elongation waiting times at individual codons (Tunney *et al.*, 2018; Duc and Song, 2018) However, our model does not explicitly account for these factors. Future extensions of the model incorporate a mixture An alternative possibility is to fit the PANSE model using elongation waiting times estimated for each codon within each gene using a machine learning framework that accounts for other upstream and downstream factors (Tunney *et al.*, 2018).

Although model comparisons support the existence of significant variation in the nonsense error rates of sense codons, most of our estimates are imprecise, spanning multiple orders of magnitude. This is not entirely unexpected given that nonsense errors are still generally rare. In contrast, our estimate assuming a uniform nonsense error rate (such that differences in NSE probabilities were only due to differences in waiting times) was precise and generally consistent with previous work (Sin *et al.*, 2016; Duc and Song, 2018). Statistical power to estimate variation in nonsense error rates across sense codons may be gained by explicitly grouping codons hypothesized to have similar NSE rates. Such a model will be implemented as part of future work.
