## Supplementary figures and images for "The Importance of Nonsense Errors: Estimating the Rate and Implications of Drop-Off Errors during Protein Synthesis"

### S1 Fig

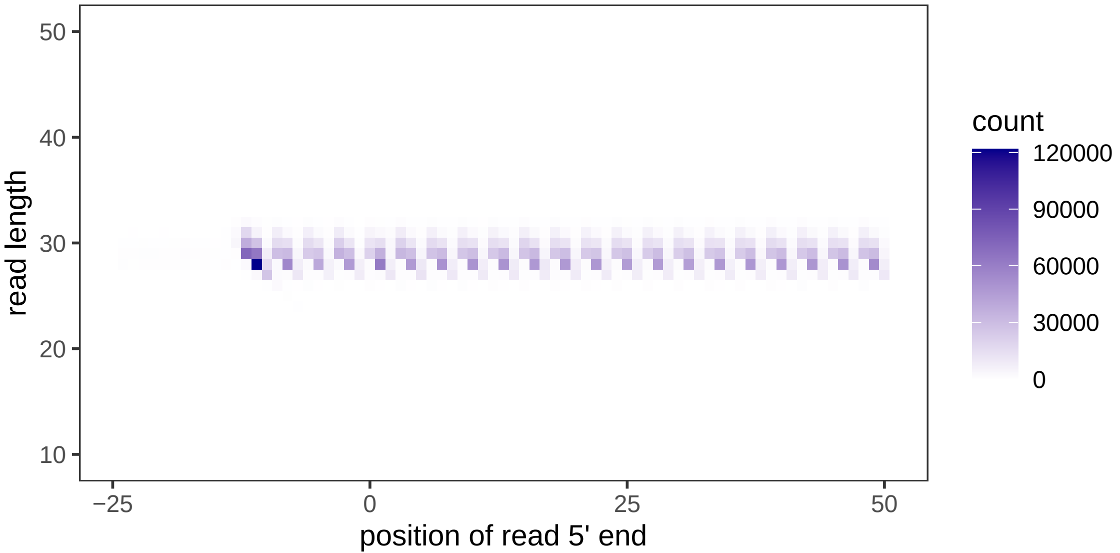

### S2 Fig

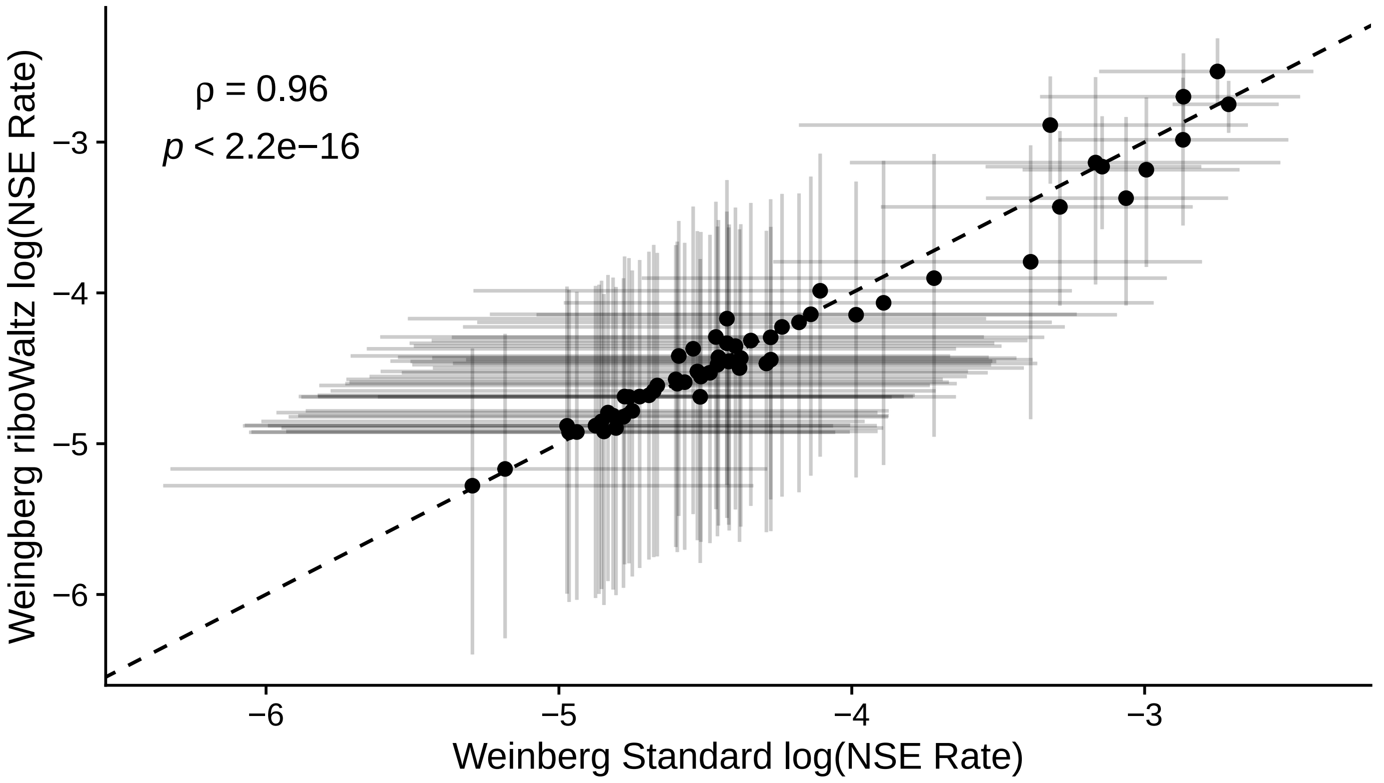

### S3 Fig

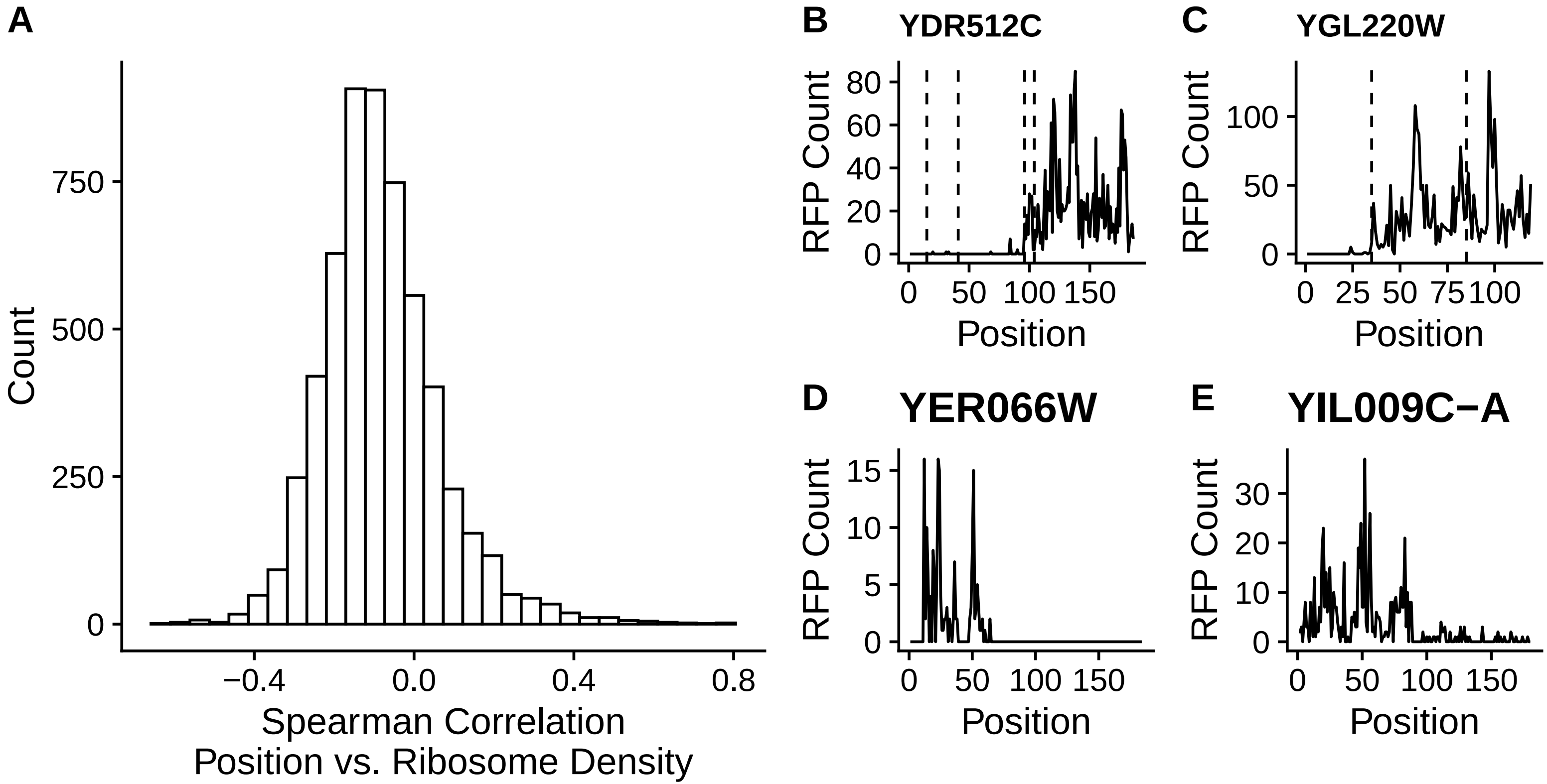

### S4 Fig

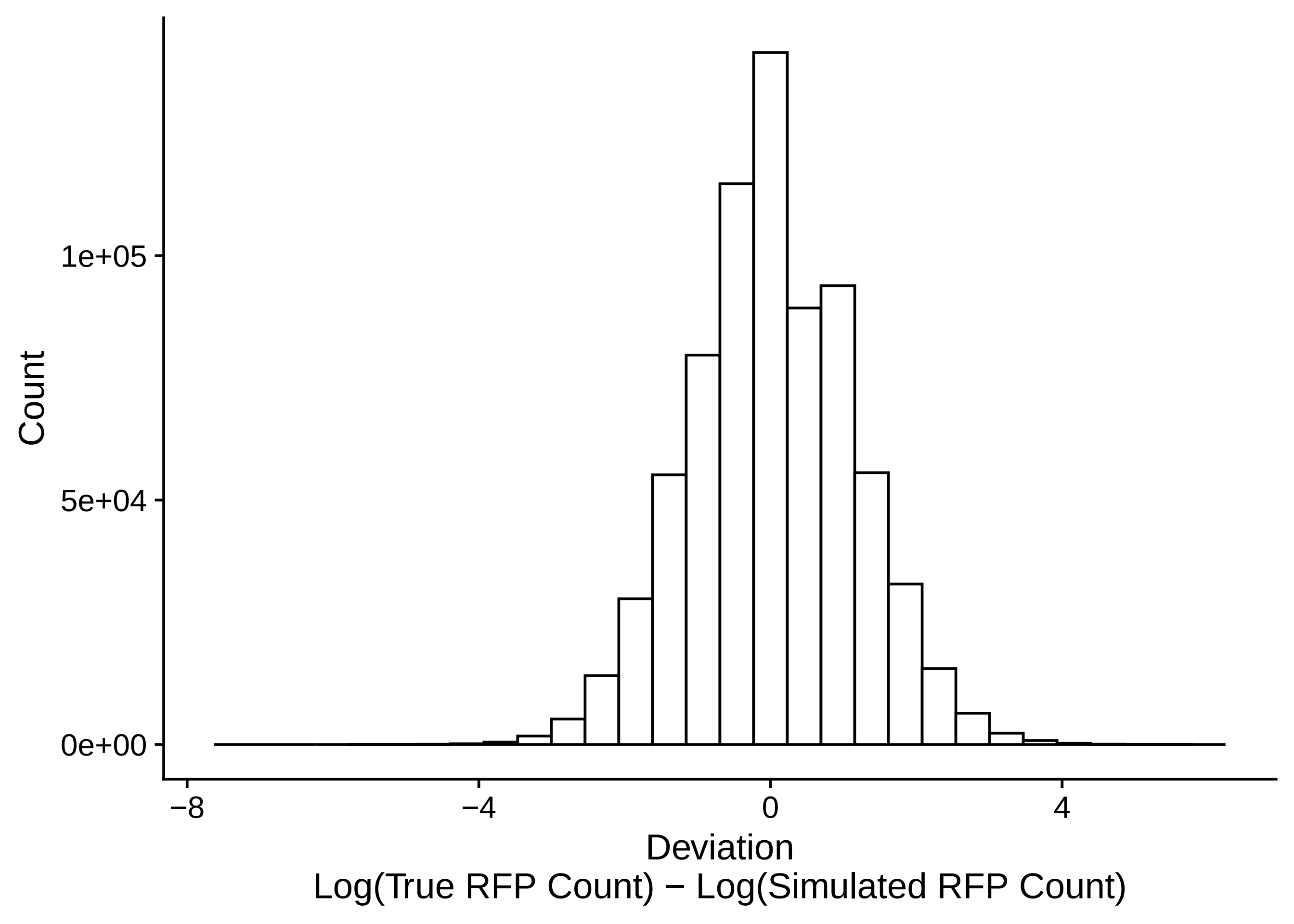

### S5 Fig

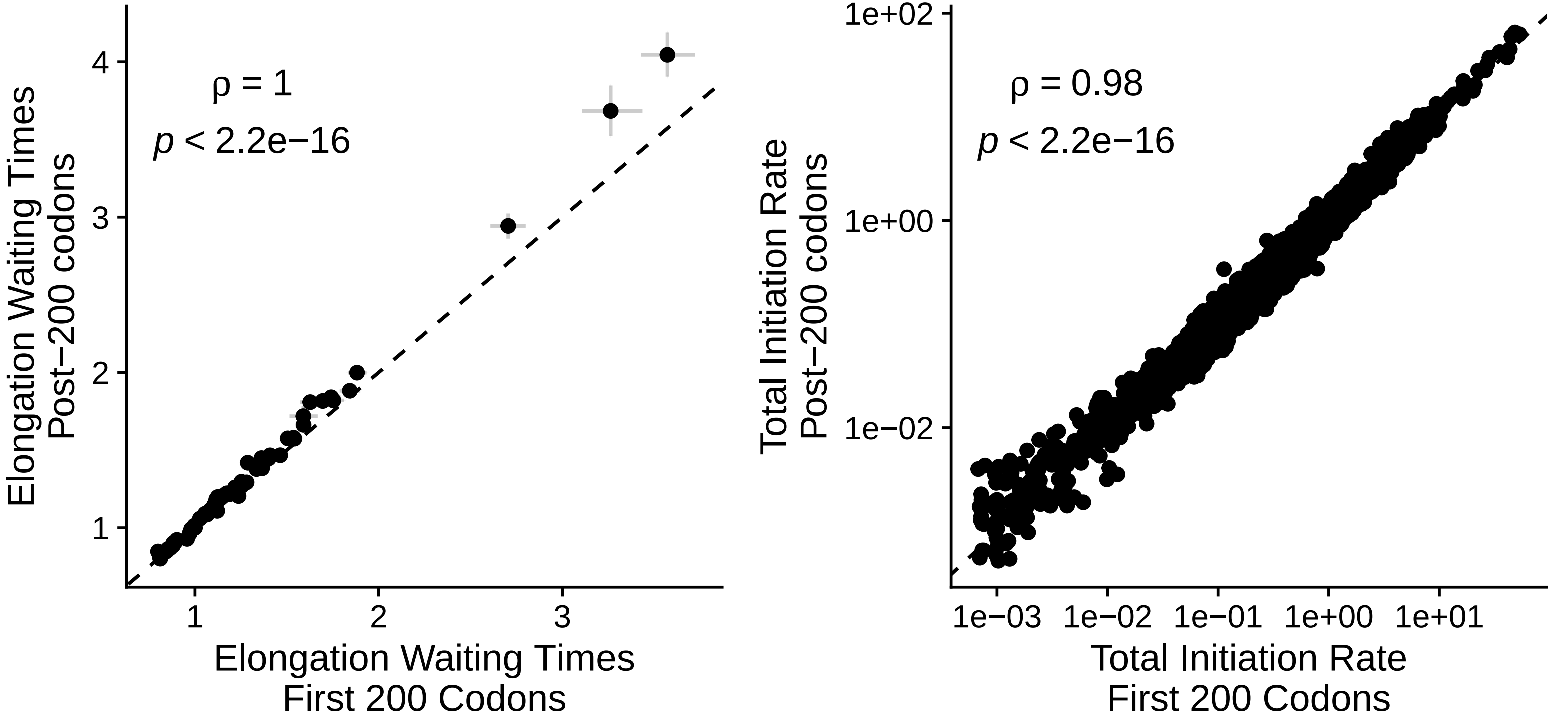

### S6 Fig

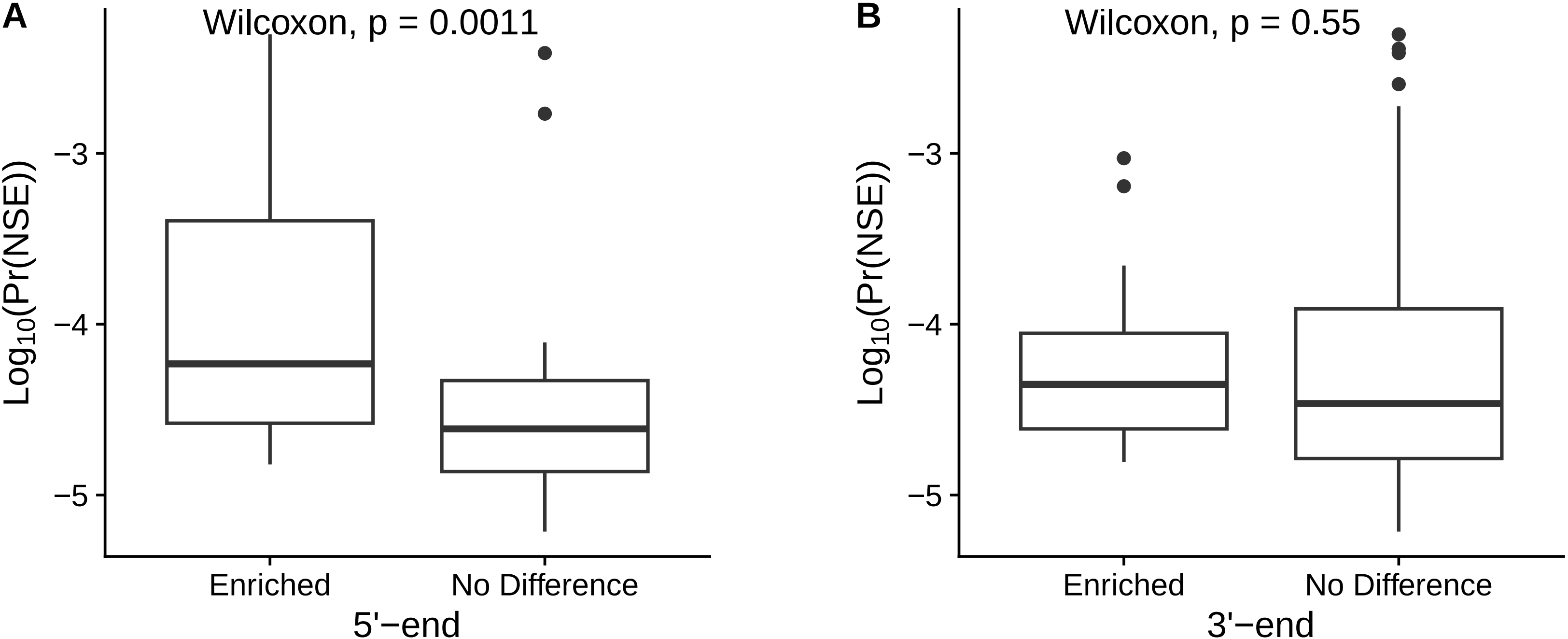

### S7 Fig

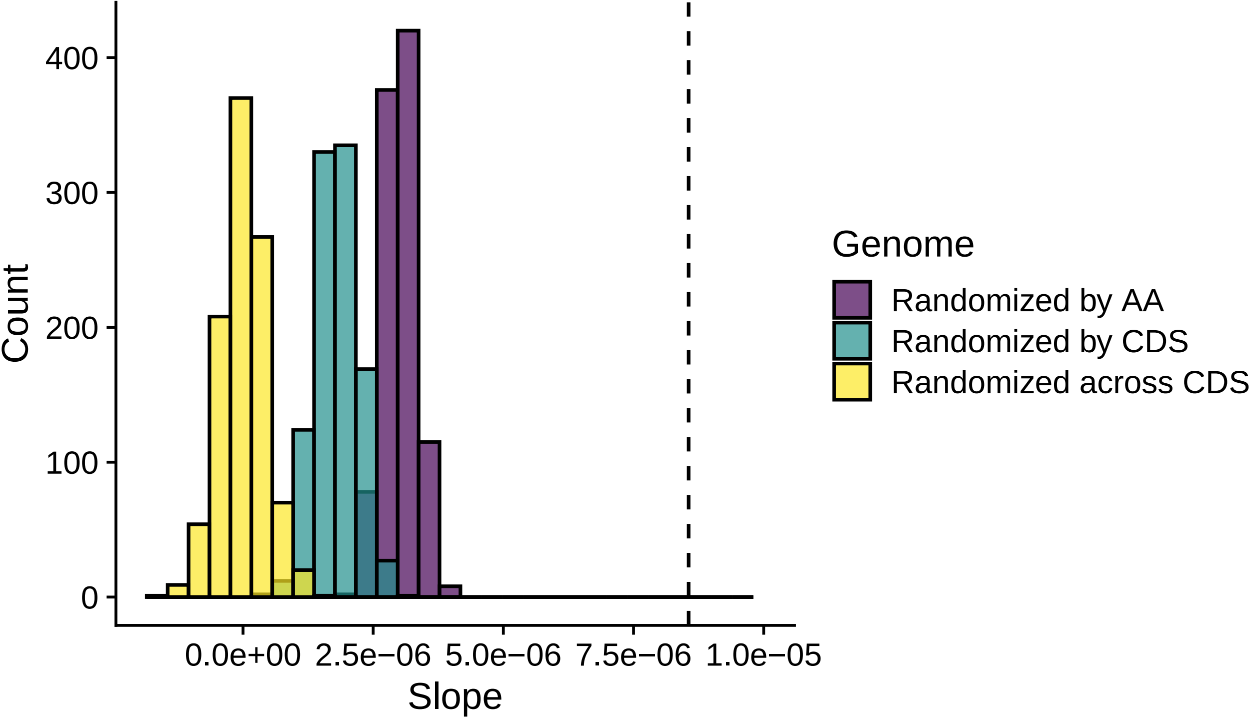

### S8 Fig

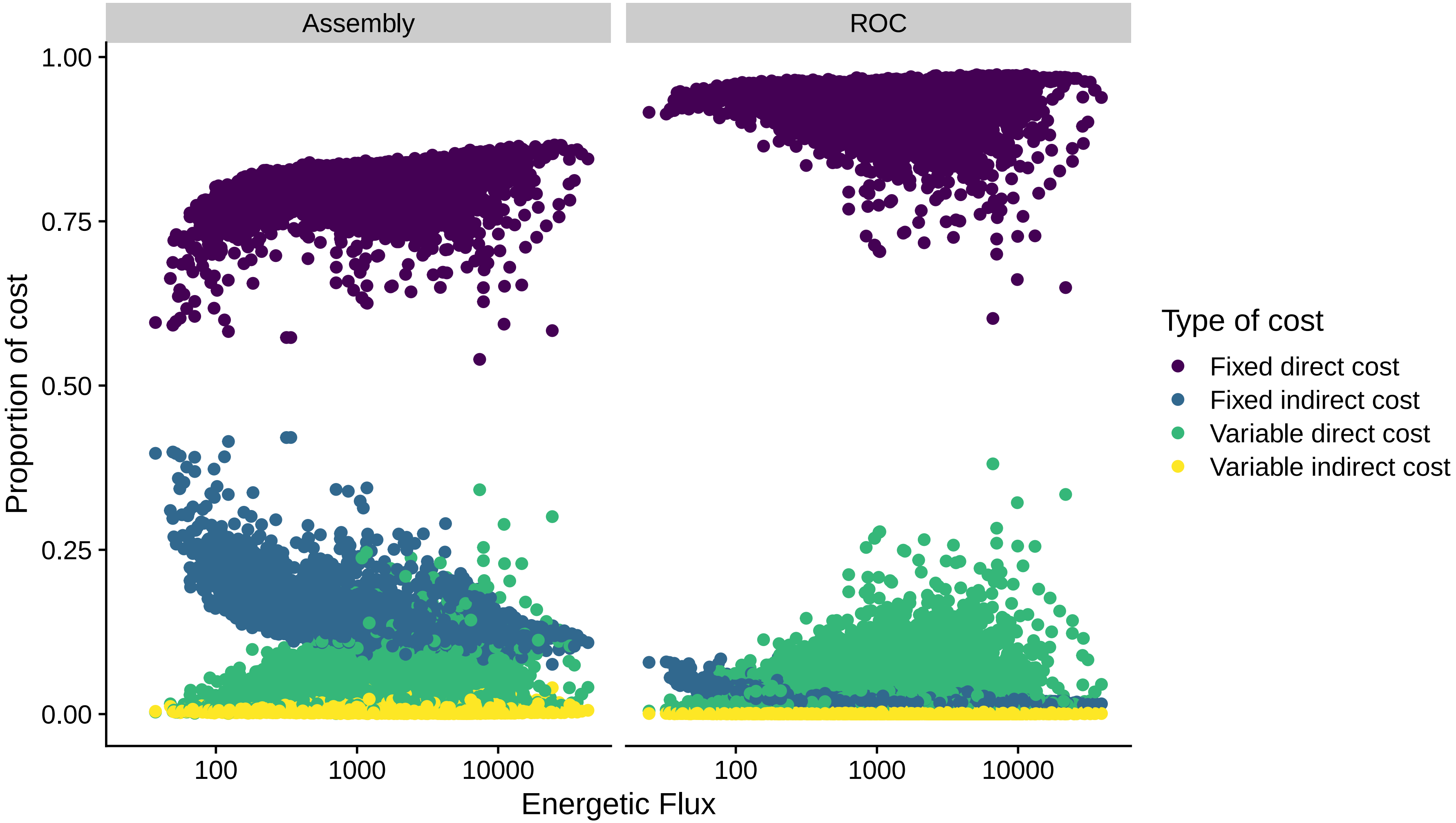

### S9 Fig

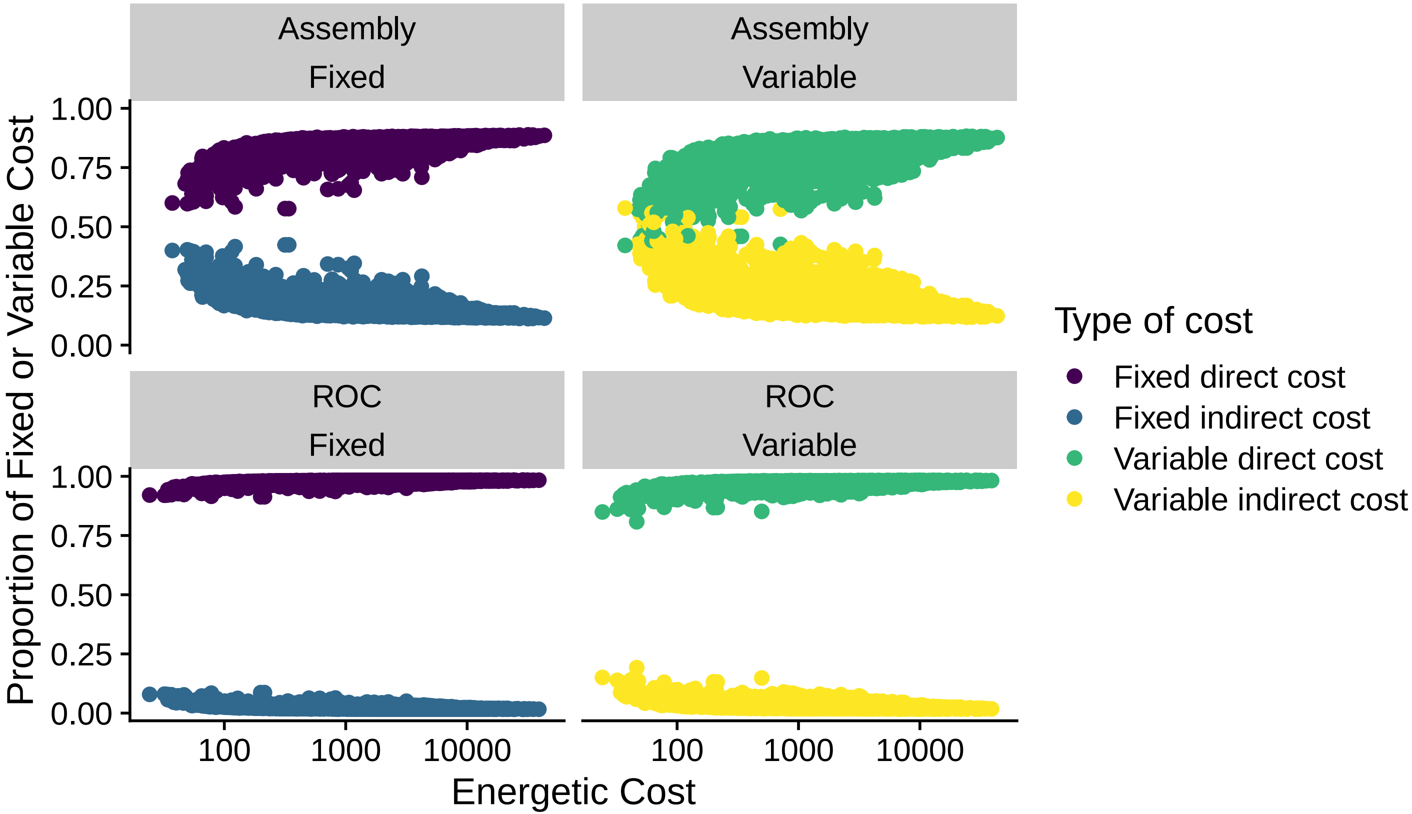

### S10 Fig

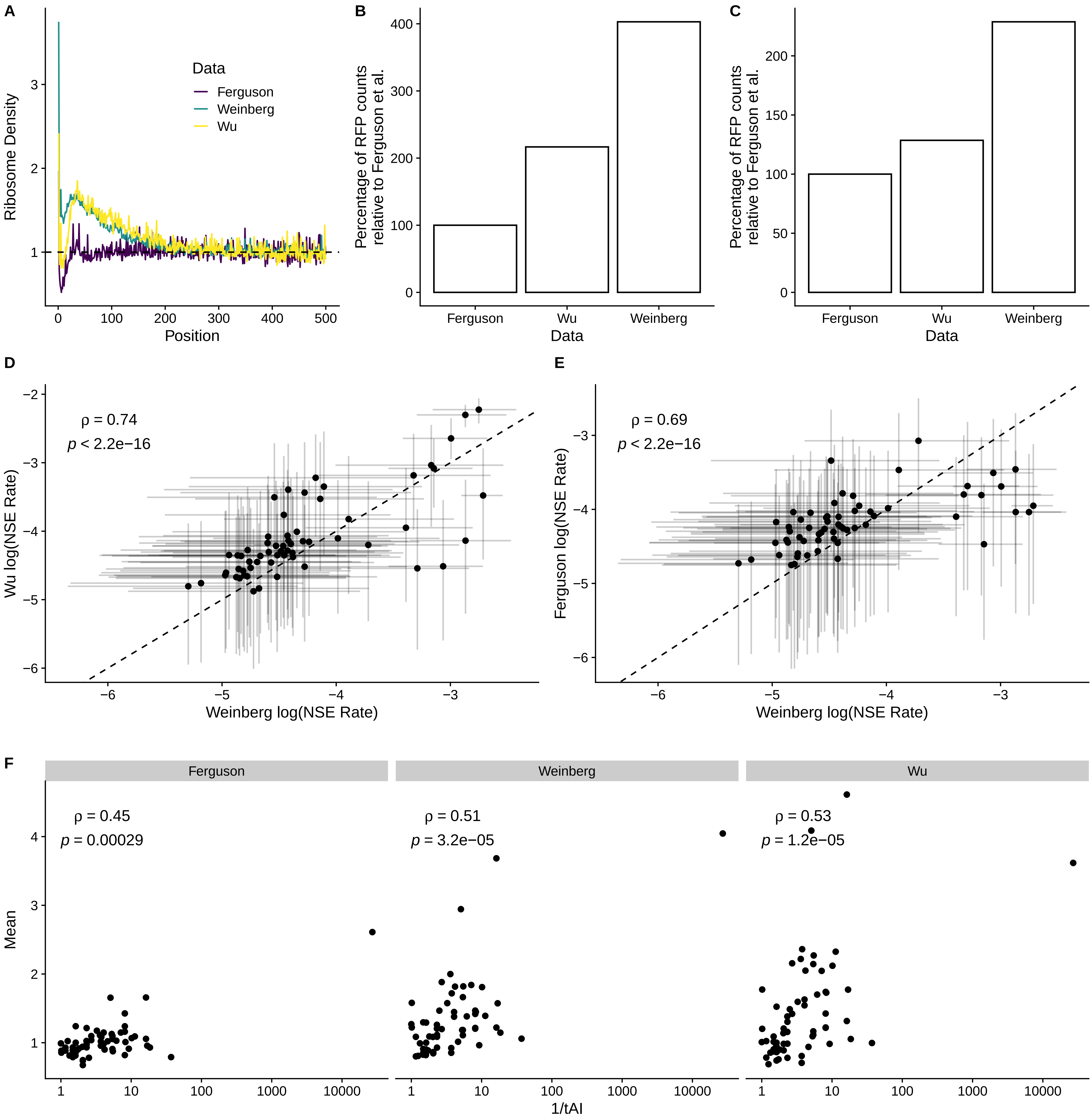
